# Metabolomics biomarkers of frailty: a longitudinal study of aging female and male mice

**DOI:** 10.1101/2025.01.22.634160

**Authors:** Dantong Zhu, Judy Z. Wu, Patrick Griffin, Brady A. Samuelson, David A. Sinclair, Alice E. Kane

## Abstract

Frailty is an age-related geriatric syndrome, for which the mechanisms remain largely unknown. We performed a longitudinal study of aging female (n = 40) and male (n = 47) C57BL/6NIA mice, measured frailty index and derived metabolomics data from plasma samples. We identify differentially abundant metabolites related to aging, determine frailty related metabolites via a machine learning approach, and generate a union set of frailty features, both in the whole cohort and in sex-stratified subgroups. Using the features, we perform an association study and build a metabolomics-based frailty clock. We find that frailty related metabolites are enriched for amino acid metabolism and metabolism of cofactors and vitamins, include ergothioneine, tryptophan, and alpha-ketoglutarate, and present sex dimorphism. We identify B vitamin metabolism related flavin adenine dinucleotide and pyridoxate as female-specific frailty biomarkers, and lipid metabolism related sphingomyelins, glycerophosphoethanolamine and glycerophosphocholine as male-specific frailty biomarkers. These associations are confirmed in a validation cohort, with ergothioneine and perfluorooctanesulfonate identified as robust frailty biomarkers. In summary, our results identify sex-specific metabolite biomarkers of frailty in aging, and shed light on potential mechanisms involved in frailty.

## Introduction

With the success of medical innovations and public health interventions, people are living much longer. However, aging is highly heterogeneous and there is extreme variability in health and function amongst different individuals of the same age^1^. Such variability in health can be captured by the concept of ’frailty’, a measurement of overall decline in health with age^2^. Frailty can be quantified using a frailty index (FI), which counts the proportion of age accumulated health-related deficits present in an individual^3,4^. Higher FI values indicate a greater degree of frailty and are associated with an increased susceptibility to diseases and mortality^4,5^. Frailty indices have been adopted for use in other mammals, including mice^6^.

Whilst frailty assessments are commonly used in both the clinic and research, there are no accepted frailty biomarkers^7^, and very little is known about the underlying molecular mechanisms of frailty, distinct from aging. Identification of frailty biomarkers would be beneficial in enabling earlier identification and tracking of frailty over time, development and testing of treatments and interventions^7^ and contribute to our understanding of the biological pathways underlying the development of frailty^8^. Metabolomics is an emerging field that enables comprehensive and quantitative metabolite assessment in biological samples. Circulating metabolites can provide a snapshot of the metabolic status of an individual, and as such have the potential to be both biomarkers, and provide insight into biological pathways changed in frailty and age.

There are known metabolic changes in aging, and in fact many of the ’hallmarks’ of aging are linked to unfavorable metabolic shifts^9^. Less is known about metabolic changes in frailty, although studies have shown that glucose intolerance and insulin dynamics are closely linked to physical frailty in both humans and mouse models^10,11^. Metabolomics studies of aging in humans are beginning to identify specific metabolite markers^12^. Elevated high- and decreased low-density lipoproteins are well established in older individuals, and associated with poor clinical outcomes^13^. Changes in amino acids are observed in aging, including increased tyrosine and decreased tryptophan^14,15^. Both lipids and amino acids are extensively related to nutrient sensing pathways, such as the mammalian target of rapamycin (mTOR)^16^ that acts as a central regulator in aging^17^. Oxidative stress and inflammation- related metabolites are also associated with aging, particularly acylcarnitines, sphingomyelins^18^, and cytochromes P450 metabolites^19^. However, the majority of these metabolomics studies are cross-sectional in design, comparing separate groups of young and old individuals, and there are few studies exploring how metabolites change longitudinally within the same individuals as they age^20,21^. Although early metabolomics studies focused on associations with chronological age only, there is a growing focus on metabolomics studies of frailty in humans, and these studies have revealed associations with energy and nutrition metabolism^22^ and with amino acid metabolism^23,24^. While these studies hint at a strong link between frailty and metabolism, they are limited by small sample sizes, and cross-sectional designs.

Additionally, sex dimorphism in aging is widely observed across many levels. Most notably, at every age, women are more frail than men, despite having longer life expectancy^25^. There are also clear sex differences in the risk and prevalence of age-related diseases, including metabolic diseases^26,27^. Many studies have revealed clear sex differences in metabolic aging across multiple tissues including blood^28^, brain^29^, and adipose tissue^30^. Sex differences in metabolites related to lipid metabolism, such as cholesterol^30^ and sex steroid hormones^30^, amino acids and acylarnitines^31^ are widely observed, and such differences can be age- dependent. The mechanisms, and especially metabolic mechanisms, underlying these sex differences in aging and frailty are not well understood, despite some recent efforts^25,32,33^. Although studies have explored baseline sex differences in circulating metabolites, as far as we are aware, there are currently no longitudinal metabolomics studies exploring sex differences in frailty.

Here, we completed a longitudinal study of female and male mice and generated matched metabolomics and frailty data across 5 time points. We use time-course and network analysis to identify age related metabolites, apply machine learning algorithms to select frailty-related metabolite features, perform an association study on frailty features, and build a metabolite frailty clock. We reveal that age-related metabolites are enriched in lipid metabolism, and suggest that amino acid metabolism and metabolism of cofactors and vitamins are enriched for frailty related metabolites. In particular, we demonstrate strong sex differences in metabolite features and their associations with frailty. We confirm these findings in a validation cohort, specifically finding consistent associations for 9 candidate frailty biomarkers, and the metabolite frailty clock achieves better prediction performance than age and sex alone, but only in male samples. Our results provide candidate metabolomic biomarkers of frailty for future testing in clinical studies, and provide insights into possible mechanisms underlying sex differences in frailty and aging.

## Results

### Metabolomics data variation

We performed a longitudinal study of female (*n* = 40) and male (*n* = 47) C57BL/6NIA mice at 5 time points, and derived metabolomics data for a total of 321 samples that have valid metabolomics data (**Table1**). In order to investigate aging- and frailty- related metabolites and mechanisms in naturally aging mice, we used non-NMN treated mice (female, *n* = 20; male, *n* = 24) as the discovery cohort for the ensuing analysis (**Fig.1**). To investigate the metabolomic data variation, we performed a principal component analysis (PCA), including a set of 781 metabolites. The PCA plot indicated clear separation of samples across time points (by PC1) and by sex (by PC2) (**Supplementary Fig.1a**). We then performed linear regression analyses on PCs and observed clustering of factors of interest (e.g., sex, time point, mouse ID) in the associations with PCs (**Supplementary Fig.1b**) based on *p*-values. We selected time points and sex as representative variables in the ensuing analysis as they showed the smallest *p*- values among the factors within the same cluster. Mouse ID also showed an association with PC2 at a significant level and was included to account for repeated measurements on the same mouse.

**Table 1.** List of female and male samples.

|  | Discovery cohort |  |  |  |  |  | Validation cohort |  |  |  |  |  |
| --- | --- | --- | --- | --- | --- | --- | --- | --- | --- | --- | --- | --- |
|  | Female<br>(n = 20) |  |  | Male<br>(n = 24) |  |  | Female<br>(n = 20) |  |  | Male<br>(n = 23) |  |  |
| Time points | Samples<sup>a</sup> | Age<sup>b</sup> | Frailty index<sup>b</sup> | Samples | Age | Frailty index | Samples | Age | Frailty index | Samples | Age | Frailty index |
| BL | 19 | 393 | 0.14 [0.12, 0.16] | 21 | 386 | 0.19 [0.18, 0.21] | 16 | 393 | 0.15 [0.13, 0.16] | 18 | 386 | 0.19 [0.18, 0.22] |
| T2 | 20 | 541 | 0.24 [0.22, 0.26] | 22 | 539 | 0.23 [0.21, 0.25] | 19 | 541 | 0.21 [0.20, 0.23] | 23 | 539 | 0.22 [0.20, 0.24] |
| T3 | 17 | 624 | 0.29 [0.26, 0.31] | 23 | 635 | 0.26 [0.24, 0.28] | 19 | 624 | 0.24 [0.21, 0.26] | 23 | 635 | 0.25 [0.23, 0.32] |
| T4 | 11 | 756 | 0.28 [0.26, 0.34] | 19 | 773 | 0.27 [0.23, 0.28] | 12 | 756 | 0.28 [0.23, 0.32] | 20 | 773 | 0.27 [0.26, 0.31] |
| T5 | NA | NA | NA | 9 | 910 | 0.37 [0.35, 0.45] | 4 | 899 | 0.33 [0.29, 0.36] | 6 | 910 | 0.44 [0.40, 0.45] |

a. Number of samples that have valid metabolomics data
b. Age (days) at assessment for frailty
c. Median [Lower quartile, Upper quartile]

**Fig.1.**
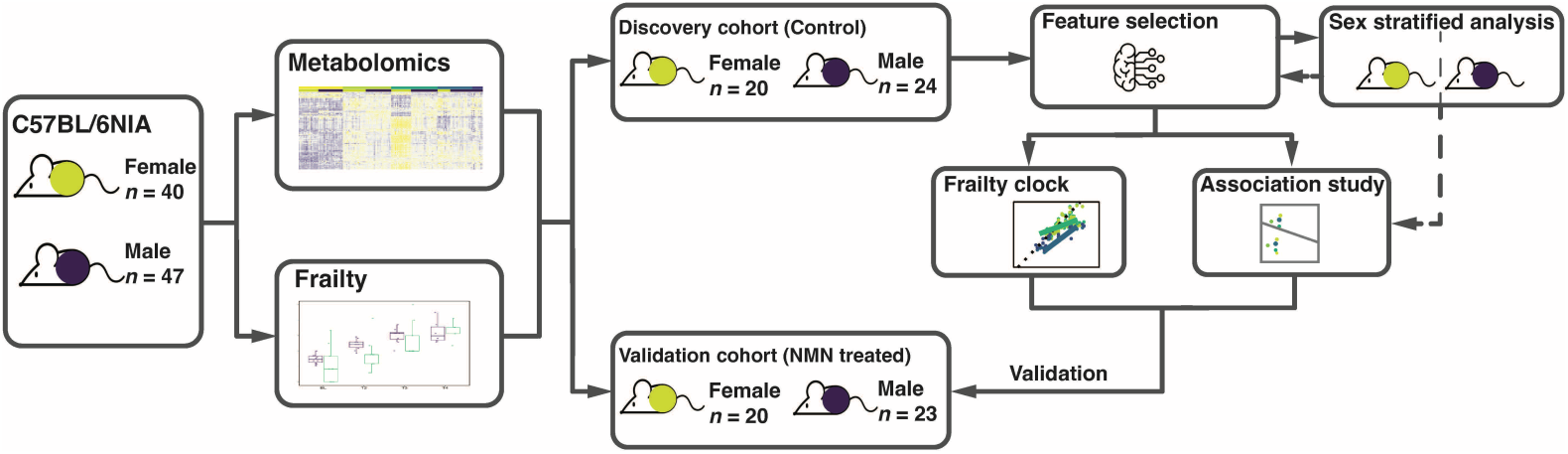
Schematic diagram of the workflow. The longitudinal study starts with female (*n* = 40, yellow circles) and male (*n* = 47, blue circles) C57BL/6NIA mice. Frailty was assessed and blood samples were collected at 5 time points from BL to T5 (exact days of experiments are shown in Table1). Plasma samples were derived from blood samples and were then subjected to metabolite quantification. In order to investigate metabolites related with natural aging and frailty, feature selection, sex stratified analysis, association study and the frailty clock were all performed in the control samples without intervention as the discovery cohort. The metabolite biomarkers and a metabolite clock for frailty were then tested in the validation cohort.

### Metabolomic signatures of aging across sexes

After the determination of covariates, we performed metabolite differential abundance analyses to identify metabolites that were related to general aging. That is, metabolite abundances that significantly changed for these mice across the sampled time points. We considered the pattern of metabolite abundance globally over time, by fitting a time series smoothing spline, accounting for mouse ID and sex. We found 527 (67.5% of total detected metabolites) differentially abundant metabolites (DAMs) over the time-course within all mice (both females and males) (**Supplementary Fig.2a**).

In order to select subsets of metabolites with similar abundance over the time course and, more importantly, highly related to aging independent of sex, we performed co-abundance network analysis on the 527 metabolites derived above. We derived two metabolite subsets, subset1 (*n* = 200) and subset2 (*n* = 125) (**Supplementary Fig.2b**) of which the eigenvalues showed strong associations (*p* < 0.001) with age, presenting a generally decreasing trend in abundance in aging (**Fig.2a**). Significantly higher proportions of metabolites within the amino acids super- pathway were observed in subset 1 (69 metabolites, 34.5% of subset1, χ^2^ (df = 2, *N* = 781) = 25.3, *p* < 0.001) and those in the lipids super-pathway for subset 2 (100 metabolites, 80% of subset2, χ ^2^ (df = 2, *N* = 781) = 141.1, *p* < 0.001), compared to the rest of metabolites.

**Fig.2.**
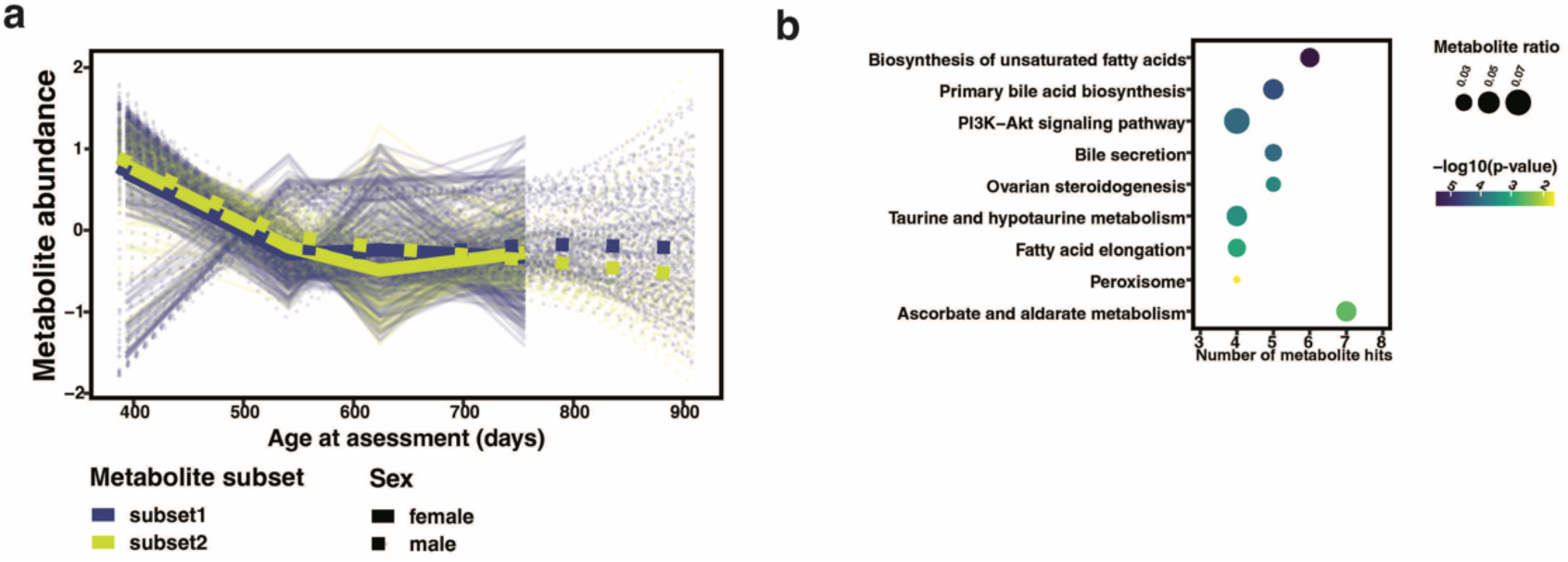
Sex independent age-related differentially abundant metabolites in longitudinal study. Differential abundance analysis was performed using all samples (excluding time point T5) in the study. Sex-independent age-related differentially abundant metabolites (DAMs) were selected from comparisons of the mixture of female and male samples at different time points and by controlling for a 5% Benjamini-Hochberg false discovery rate (adjusted *p*-values < 0.05). These DAMs were then subjected to a co-abundance analysis, and subset1 and subset2 were determined to be significantly associated with age by linear mixed models. **(a)** Dynamics of metabolite abundance in each sex, derived from two subsets (subset1, n = 200 metabolites; subset2, n = 125). After determining the hub metabolites based on metabolite correlation with age and module membership, hub metabolites were subjected to metabolite set enrichment analysis. **(b)** Over-represented pathways (y-axis) from the hub metabolites from the two subsets. The number of hits (metabolite) from the hub metabolites set is shown by x-axis, ratio of the hit number to total metabolites in the enriched pathway is represented by dot size and p- value is colored by levels.

Metabolite set enrichment analysis results on two subsets aligned with the above classification, with amino acid metabolism (subset1) and lipid metabolism (subset2) pathways over- represented (**Supplementary Fig.2c** and **d**). To further select metabolites that play important roles in aging, we selected 86 hub metabolites, 46 metabolites from subset1 and 40 from subset2, based on module membership in the network and significance (correlation coefficient between eigenvalue and age) (**Supplementary Fig.2e**). These 86 metabolites were defined as core age-related metabolites in the ensuing analyses, and include guanidinoacetate, methylamalonate (MMA) and sphingomyelin species (**Supplementary Table1**). 54.7% of these metabolites (47 total, 9 from subset1 and 38 from subset2) are from the lipid super-pathway, also evidenced by enrichment analysis (**Fig.2b**). This result suggests lipid metabolism is among the key mechanisms contributing to general aging.

### Sex specific metabolomic signatures of aging

To identify sex specific metabolomic signatures, we investigated DAMs within 1) females only (significant change in abundance in the whole time frame), n = 498 DAMs (63.8%, 498/781), 2) males only, n = 253 DAMs (32.4%, 253/781); and 3) sex differences (significantly differentially abundant in females and males considering the whole time frame), n = 331 DAMs, (42.4%, 331/781). The results suggest significant sex differences in metabolite abundance in the aging process.

It was interesting to observe a common set of 97 metabolites after merging the above three sets of DAMs with the 527 DAMs derived from the mixture of both sexes (sex-independent) (**Fig.3a**; **Supplementary Table2**). These metabolites not only were related to aging in both sexes, but also presented sex differences in aging (**Supplementary** Fig.3). Notably, these included 8 acylcarnitines, for instance, oleoylcarnitine (C18:1) and palmitoleoylcarnitine (C16:1). Among the 97 metabolites, 41 are in lipid and 22 in amino acid super pathways, representing 64.9% of the 97 metabolites. Enrichment analysis revealed 11 KEGG pathways overrepresented (**Fig.3b**), mostly within lipid metabolism and digestive system pathways.

**Fig.3.**
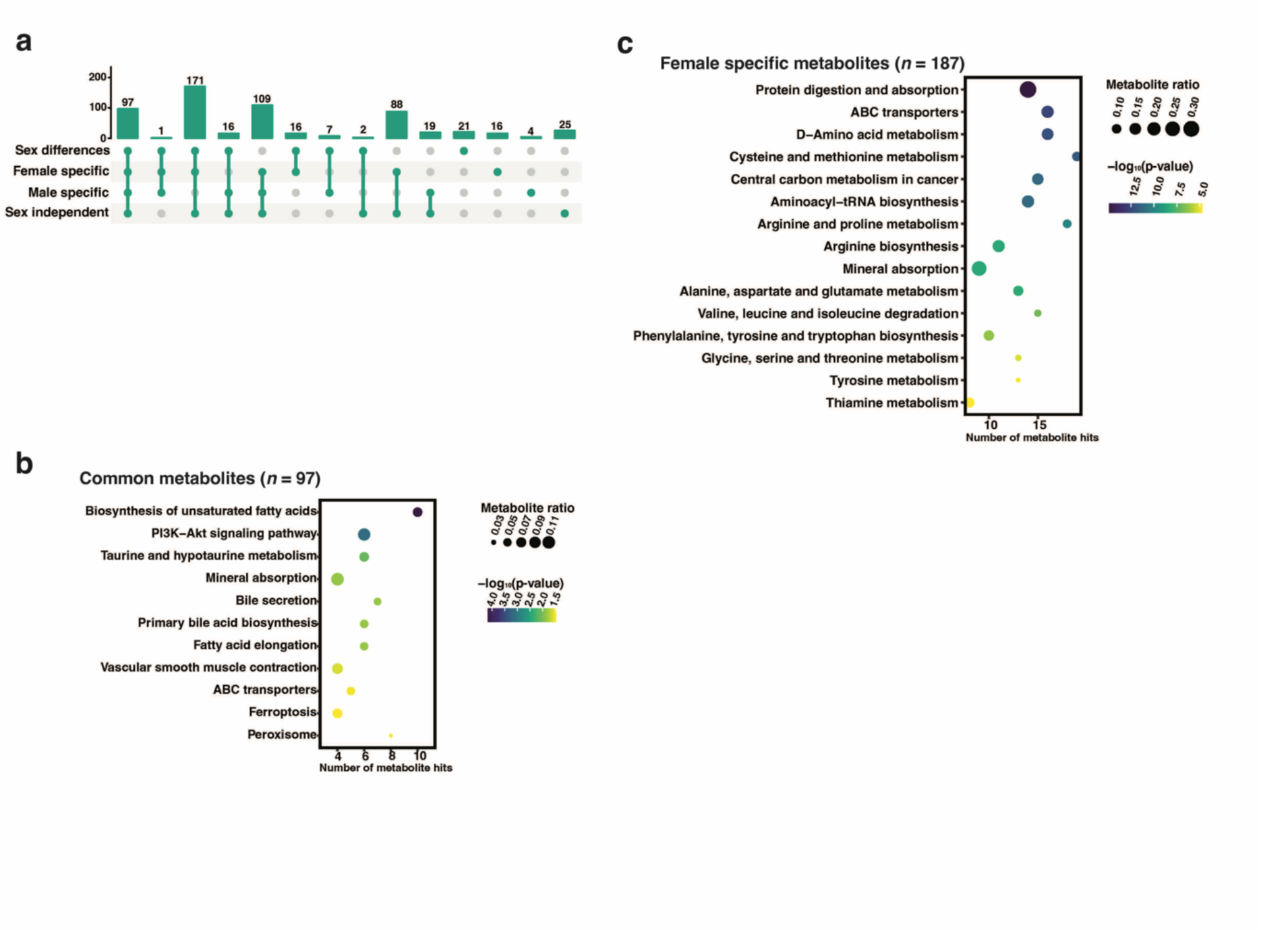
Comparisons of differentially abundant metabolites determined in four groups. Differential abundance analysis was performed using all samples (excluding time point T5) in the study. Age-related differentially abundant metabolites (DAMs) were determined by comparisons within four groups: the mixture of females and males (sex independent), female specific, male specific, and sex differences, and by controlling for a 5% Benjamini-Hochberg false discovery rate (adjusted *p*-values < 0.05). **(a)** UpSet plot showing the common DAMs derived from the comparisons. **(b)** Over-represented pathways (y-axis) from the 97 common metabolites of four groups by metabolite set enrichment analysis. **(c)** Over-represented pathways (y-axis) from the 187 female specific metabolites markers that also present sex differences. The number of hits (metabolite) from the hub metabolites set is shown by x-axis, ratio of the hit number to total metabolites in the enriched pathway is represented by dot size and p-value is colored by levels.

Apart from the common set, among the 331 metabolites that are different between males and females across the investigated timeframe, there are only 21 metabolites that present sex differences (distinct abundance in females and males) and do not present significant abundance differences over time (**Supplementary Table2**). This indicates that the majority of sex differential metabolites also change with age. In terms of female-specific metabolites that are changed with age, that is, metabolites detected in both female and sex difference DAM sets, the 187 in this category included amino acids and acylcarnitines and were enriched for amino acid metabolism pathways (**Fig.3c**; **Supplementary Table2**). Many fewer male-specific age-related metabolites were observed, with a total of only 23, including phosphocholine and spermine. One metabolite, corticosterone, changed with age in both females and males separately (**Supplementary** Fig.4), but was not detected to change with age when the entire cohort of mice was considered. Altogether, we found metabolites involved in lipid metabolism and digestive system pathways contribute to aging and present strong sex differences. Specifically, amino acid metabolism-related metabolites are associated with aging in female mice.

### Sex independent metabolite features of frailty

Having identified age-related metabolites, we were interested to identify metabolites specifically associated with frailty. Frailty is a complex geriatric syndrome. For each mouse at a certain time point, FI is composed of the base FI (median FI of the corresponding sex and age group) and devFI (the deviation of individual FI from the median FI at corresponding age- and sex-specific group). By definition, base FI is highly related to age, but devFI is age independent (**Supplementary** Fig.5).

In order to find metabolites related to frailty, we performed feature selection outlined in **Fig.4a**. We investigated metabolites that were related to both FI and devFI, by performing feature selection with elastic net regularization, via a 100 times repeated 5-fold cross validation approach. Based on the rank of presence frequency, we selected 156 and 149 metabolites predictive of FI and devFI, respectively (**Supplementary Fig.6a**; **Supplementary Table3**). 86 of these metabolites were identified as both devFI and FI features (**Fig.4b**), suggesting both overlapping and distinct metabolite signatures of FI and devFI. The majority of identified FI and devFI metabolites were within the amino acid and lipids super pathways (**Supplementary Fig.6b**). Three metabolites were simultaneously identified as FI-, devFI- and age-metabolites, including ergothioneine that decreases with age and frailty in both females and males (**Supplementary** Fig.7). When looking at the top enriched KEGG pathways for FI and devFI, there were 11 common pathways (out of the top 15 by p-value) across both groups (**Supplementary Fig.6c** and **d**), including 7 amino acid metabolism pathways, nicotinate and nicotinamide metabolism, pantothenate and CoA biosynthesis, pyruvate metabolism and ABC transporters. To further identify core-metabolites related to frailty, we derived 21 FI-age and 86 devFI features by merging age-related metabolites (86 hub metabolites) and devFI metabolites with the FI metabolites respectively, resulting in a set of 104 union features (**Fig.4b**) which are enriched for amino acid metabolism and metabolism of cofactors and vitamins pathways (**Fig.4c**). These results suggest these metabolic pathways, notably, amino acids and B vitamin metabolism are specifically important in the development of frailty.

**Fig.4.**
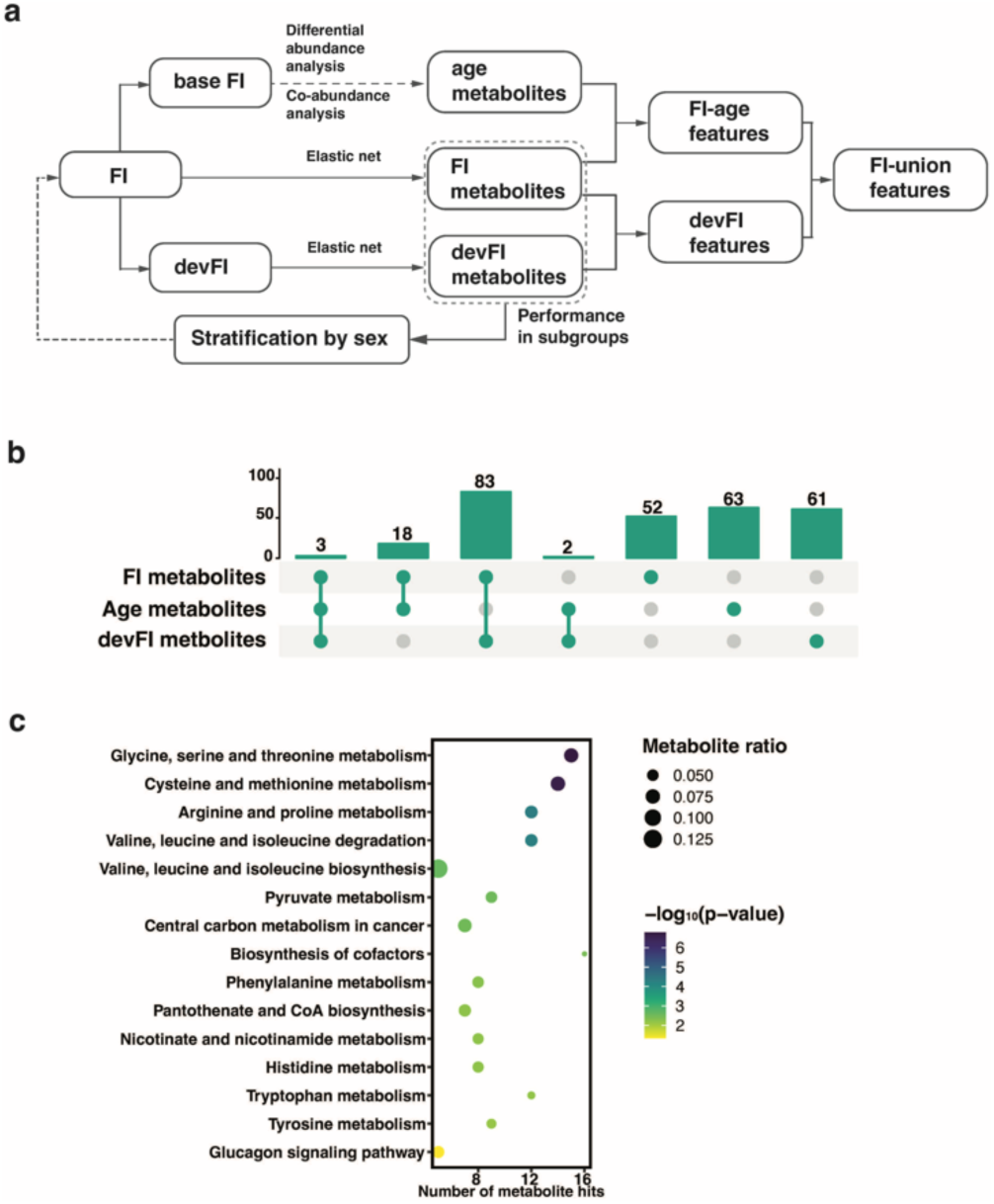
Selection of frailty related features **(a)** Schematic diagram for the workflow of the feature selection. Frailty index (FI) is composed of base FI and devFI (deviation from the age- and sex- group median FI). Base FI is age related, hence leads to age metabolites. FI and devFI metabolites are derived from elastic net regularization regression via a 100 times repeated 5-fold cross validation approach. FI metabolites are merged with age metabolites into FI-age features and with devFI metabolites into devFI features. The FI-union features are the union of FI-age and devFI features. The workflow is performed in the whole cohort, as well as females and males after the stratification by sex. **(b)** UpSet plot showing the overlapping metabolite features from the FI-, age- and devFI- metabolites. **(c)** Over-represented pathways (y-axis) from the 104 FI-union features from the whole cohort. The number of hits (metabolite) from the hub metabolites set is shown by x-axis, ratio of the hit number to total metabolites in the enriched pathway is represented by dot size and p-value is colored by levels.

Despite the common metabolites, we observed 61 metabolites that are unique to devFI (**Fig.4b**), including hippurate, choline, hypotaurine, phenylacetyltaurine, and adenosine 5’- diphosphoribose. Enrichment analysis based on these metabolites led to efferocytosis and ABC transporters. The results suggest these metabolites and pathways are associated with frailty in a completely age- and sex-independent way.

### Association study of metabolite features with frailty outcomes

To test the associations of the individual metabolite features with frailty outcomes (**Fig.5a**), we applied linear mixed regression models and subjected the 104 union frailty features to a longitudinal association study. First, we considered only metabolite abundance at the current timepoint (Age_c_). For the current FI (FI_c_) and devFI (devFI_c_), we found 47 and 16 metabolites, respectively presented coefficients significantly different from 0 (**Supplementary Table4**).

**Fig.5.**
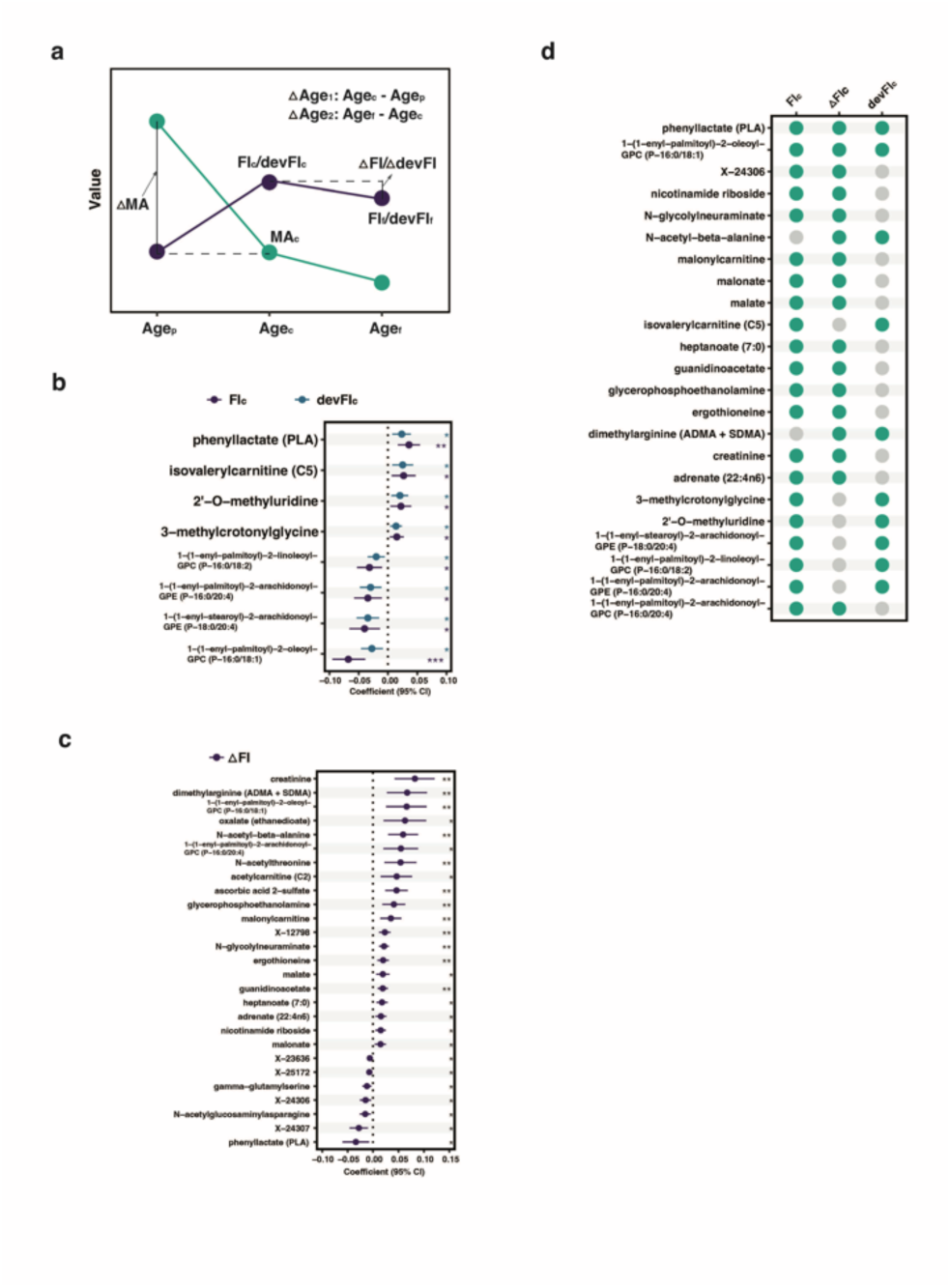
Association study in the whole cohort **(a)** Schematic diagram showing the dependent and independent variables in the linear mixed models for the association study. Dependent variables include Frailty Index (FI_c_) and devFI (devFI_c_, deviation from median FI of the age- and sex- specific group) at the current age (age_c_), FI and devFI (FI_f_/devFI_f_) at a future age (age_f_), and FI/devFI change from age_c_ to age_f_ (ᐃFI/ᐃdevFI). Independent variables include current abundance of metabolites (MA_c_), abundance change from a previous age (age_p_) to age_c_, ᐃage_1_ and ᐃage_2_. For each frailty outcome, FI-union features identified were individually subjected to linear mixed models. **(b)** Coefficients of eight metabolites of which MA_c_ presents significance in the association with both FI_c_ and devFI_c_. Metabolites are arranged by coefficients (represented by dots) for FI_c_ in descending order. The line represents the 95% confidence interval of each coefficient. **(c)** 27 metabolites of which the current metabolite abundance presents significance in the association with ᐃFI. Metabolites are arranged by coefficients (represented by dots) in descending order. Significance was determined by adjusted *p*-values via Benjamini-Hochberg false discovery rate procedure at a cutoff of 0.05, with * for *p* < 0.05, ** for *p* < 0.01, and, *** for *p* < 0.001. **(d)** List of 23 metabolites that show occurrences greater than or equal to 2. That is, MA/ᐃMA of metabolite presents significance in the association with frailty outcomes of the column.

Among the 47 FI_c_ metabolites, three metabolites (leucine, N-acetylthreonine, and X-25422), presented a significant metabolite abundance by age interaction term (**Supplementary Table5**), indicating that the association of these metabolites with FI_c_ is age-dependent. Despite this, leucine showed a generally consistent positive correlation with FI_c_ at each age group, but this was not the case for N-acetylthreonine and X-25422 (**Supplementary Fig.8a**). The remaining 45 metabolites (those without significant interaction terms, plus leucine) are associated with FI_c_ independent of age. That is, individual mice with higher abundance of these 45 metabolites are either more (19 metabolites, β > 0) or less (26 metabolites, β < 0) frail in a cohort. For devFI_c_, individual metabolites were also associated with both higher (9 metabolites, β > 0) and lower (7 metabolites, β < 0) frailty scores. Eight metabolites were identified as significantly associated with both FI and devFI (**Fig.5b**). For each of these metabolites, the coefficient of association was in the same direction, indicating the same trend of association with both FI and devFI.

Given the longitudinal nature of our dataset we were interested to observe whether metabolite abundance at any specific timepoint was associated with frailty at a future timepoint (**Fig.5a**). Unfortunately, we didn’t observe any metabolites that showed an overall significant association with future FI (FI_f_) or future devFI (devFI_f_). When focusing only on the abundance of metabolites at the baseline time point (∼400 days), we found a single metabolite, alpha-ketoglutarate, was negatively associated with both FI_f_ and devFI_f_ (**Supplementary Fig.8b**). Next, we considered whether there were associations between current metabolite abundances, and a change in FI from one timepoint to the next (ᐃFI or ᐃdevFI, **Fig.5a**). We saw no associations with ᐃdevFI but found 27 metabolites that showed significant associations with ᐃFI (**Fig.5c** and **Supplementary Table4**). No significant interaction terms were observed for these metabolites, indicating that these associations were not age-dependent. 20 metabolites (β > 0, e.g. creatinine) were associated with increased frailty and the remaining 7 (β < 0, e.g. phenyllactate) were associated with decreased frailty (**Supplementary Fig.8c**). Finally, we considered whether changing abundances of a metabolite over time (ᐃMA), might be associated with frailty (Fig.5b) but found no significant associations.

Combining the 3 sets of metabolites above (those significantly associated with FI_c_, ᐃFI, or devFI_c_) gives a total of 63 metabolites, of which 23 are present in 2 or more sets (**Fig.5d**). These metabolites represent candidate biomarkers for frailty and include phenyllactate, ergothioneine, nicotinamide riboside, creatinine, alpha-ketoglutarate, isoleucine and valine.

### Sex specific metabolite features of frailty

Sex differences are common in aging and frailty. We investigated the performance of the generalized linear models that were trained to predict FI or devFI in the whole cohort (**Fig.4a**), in the females and males separately. We found significant differences (two-tailed t-test, *p* < 0.001) between the R-squared values derived from female and male samples (**Supplementary Fig.9a**), suggesting the associations of metabolites with FI and devFI are sex specific and stratification by sex is appropriate for this analysis. Interestingly, the model performance was better in the females than males.

In order to select sex specific metabolites related to frailty, we stratified the whole cohort into female and male subgroups and re-selected metabolite features as above. For females, we derived 133 and 45 metabolites related to FI and devFI, and for males, we obtained 32 and 92, respectively. Of these only 7 were associated with FI, and 8 with devFI, in both sexes. Despite this, for both males and females, the majority of the identified metabolites were within amino acids and lipids super pathways (**Supplementary Fig.9b**), and the enriched pathways were similar between males and females. They predominantly included amino acid metabolism, metabolism of cofactors and vitamins, mineral absorption, and protein digestion and absorption related to the digestive system (**Supplementary Fig.9c** and **d**).

Following the union feature workflow (**Fig.4a**), we obtained 58 union features for females and 21 for males related to overall FI. Within the female union features, 50% of the metabolites were related to amino acid and lipid pathways, whilst the male union features were enriched in lipid super pathways (χ^2^(df = 2, *N* = 781) = 4.11, *p*-value = 0.042). Excluding the union features associated with frailty in the whole cohort, we identified 25 and 9 metabolites that are unique metabolite features identified only in females or males (**Fig.6a**). These sex specific features include kynurenate and quinolinate for females and sphingomyelin and creatinine for males. These results suggest sex specific biomarkers for frailty may be appropriate.

**Fig.6.**
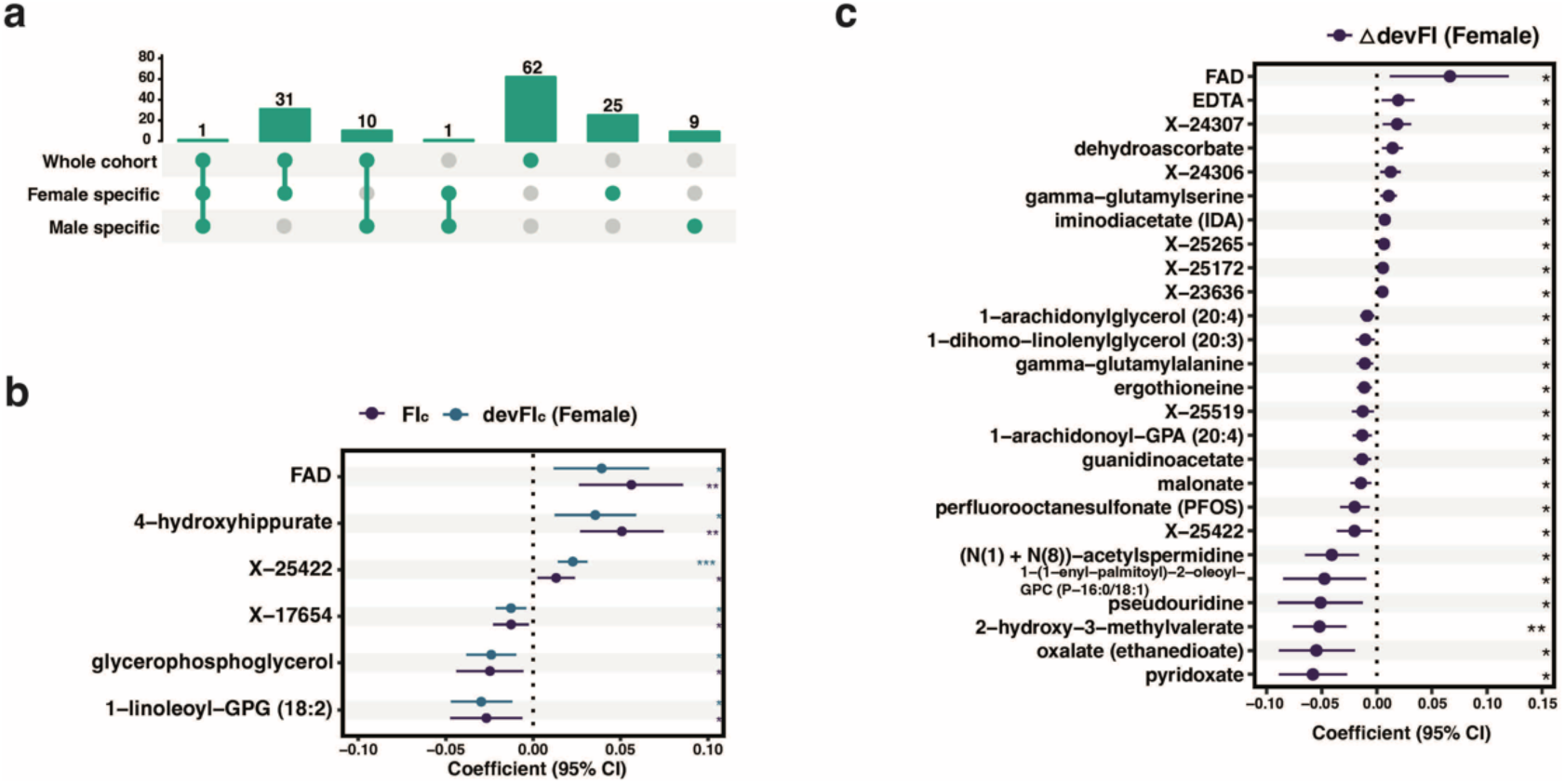
Sex independent metabolite features and association **(a)** UpSet plot showing the overlapping metabolites of frailty index union features of the whole cohort, females and males. **(b)** Coefficients of six metabolites of which metabolite change presents significance in the association with both FI_c_ and devFI_c_ in females. Metabolites are arranged by coefficients (represented by dots) for FI_c_ in descending order. **(c)** Coefficients of 26 metabolites of which metabolite change presents significance in the association with devFI change in females. The significance was determined by adjusted *p*-values via Benjamini- Hochberg false discovery rate procedure at a cutoff of 0.05, with * for *p* < 0.05, ** for *p* < 0.01, and, *** for *p* < 0.001. Metabolites are arranged by coefficients (dots) in descending order. The line represents the 95% confidence interval of each coefficient.

### Association study of sex specific frailty features

To investigate the association of individual metabolites with frailty in each sex, we performed mixed linear model regressions using the FI union features for females and males separately, as above. In females, we first considered solely the current metabolite abundance and found that 38 and 16 metabolites, respectively, were significantly associated with FI_c_ and devFI_c_ (**Supplementary Table6**). As with the whole cohort, metabolites were both positively (21 for FI_c_ and 11 for devFI_c_) and negatively (17 for FI_c_ and 5 for devFI_c_) associated with frailty outcomes. Notably, 6 metabolites were identified as both FI_c_ and devFI_c_ related (**Fig.6b**). When considering associations between current metabolite levels and either future frailty, or changing frailty levels (**Fig.5a**), we found 26 metabolites were associated with ᐃdevFI (**Supplementary Table6**).

These associations were independent of age, suggesting a relationship between metabolite levels and either increasing (10 metabolites, β > 0) or decreasing (16 metabolites, β < 0) rate of development of frailty. (**Fig.6c**). Next, we considered the relationship between frailty outcomes and changing metabolite abundances over time, and found one metabolite, ergothioneine, which was significantly associated with devFI_c_. We combined the four datasets from these female-specific frailty association studies to identify a total of 52 metabolites, of which 3 metabolites were present across 3 lists, including FAD and ergothioneine, and 23 metabolites were present across 2 lists (**Supplementary Fig.10a**). These 26 metabolites are potential female specific frailty biomarkers.

In male samples, we followed the same analysis flow. Considering current metabolite abundance, we found 19 and 12 metabolites that were associated with FI_c_ and devFI_c_ (**Supplementary Table6**). Of these, 11 metabolites were excluded as they had significant interaction terms, indicating that the association of the metabolite with FI_c_ depends on age (**Supplementary Table7**). No significance was found for the remaining regressions for the male samples. The results include three GPEs, one GPC, creatine, and phenyllactate, that may be potential male specific frailty biomarkers (**Supplementary Fig.10b**).

### Validation of the frailty associated metabolites

In order to validate the metabolites associated with frailty in an external cohort, we used female (*n* = 20) and male (*n* = 23) mice samples under long-term NMN treatment (**Table1**). To investigate if the associations of our identified frailty features with frailty outcomes persist under the intervention, we used union features identified from the whole cohort, and females and males separately, and performed the same association analysis. For sex-independent features, we found one metabolite, perfluorooctanesulfonate, that was significantly associated with current frailty (FI_c_) in the validation cohort. There were seven metabolites associated with FI change over time (ᐃFI, **Fig.5a**), including ergothioneine, guanidinoacetate, N-glycolylneuraminate, X-12798, creatinine, dimethylarginine (ADMA + SDMA), and N-acetyl-beta-alanine. In male samples only, 2-hydroxydecanoate maintained a significant association with FI_c_ at one time point (**Supplementary** Fig.11). Although NMN treatment delays frailty ^34^, the persistent association of these metabolites with frailty outcomes reveals evidence for their robustness as possible frailty biomarkers.

### Development of a metabolite-based frailty clock

To build a model to accurately predict frailty using metabolomics features in aging mice, we fit a random forest model in the discovery cohort, with FI as the dependent variable and the combination of identified union features from the whole cohort, female- and male- specific analysis (total *n = 139* metabolites) as the independent variables. We further determined the top 63 metabolite features ranked by the presence frequency (**Supplementary Fig.6a**) gave the best performance in predicting FI (**Supplementary** Fig.12). Among the 63 metabolite features that can be found in the whole cohort derived features, 24 metabolites were also identified as female-specific frailty related metabolites, and 4 as male-specific. Our final model, metabolite frailty clock, included these 63 informativity-based metabolites, age and sex. We also fit a model using all 781 metabolites detected in our study, as a comparison. Both random forest models

performed well in the discovery cohort (*R^2^ =* 0.95, RMSE = 0.022 for RF with 781 metabolites; *R^2^ =* 0.96, RMSE = 0.019 for metabolite frailty clock), and outperformed a benchmark model of merely age+sex (*R^2^* = 0.51, *RMSE* = 0.053) (**Fig.7a**). Importantly, we examined the performance of the metabolic frailty clock in the validation cohort, and although it achieves similar performance as the age+sex model trained in the validation cohort across the entire cohort (**Fig.7b**), it outperforms age+sex in male samples (**Fig.7c**). Despite the fact that there is clearly room for improvement, these results suggest that frailty can be accurately predicted in aging mice using metabolite features.

**Fig.7.**
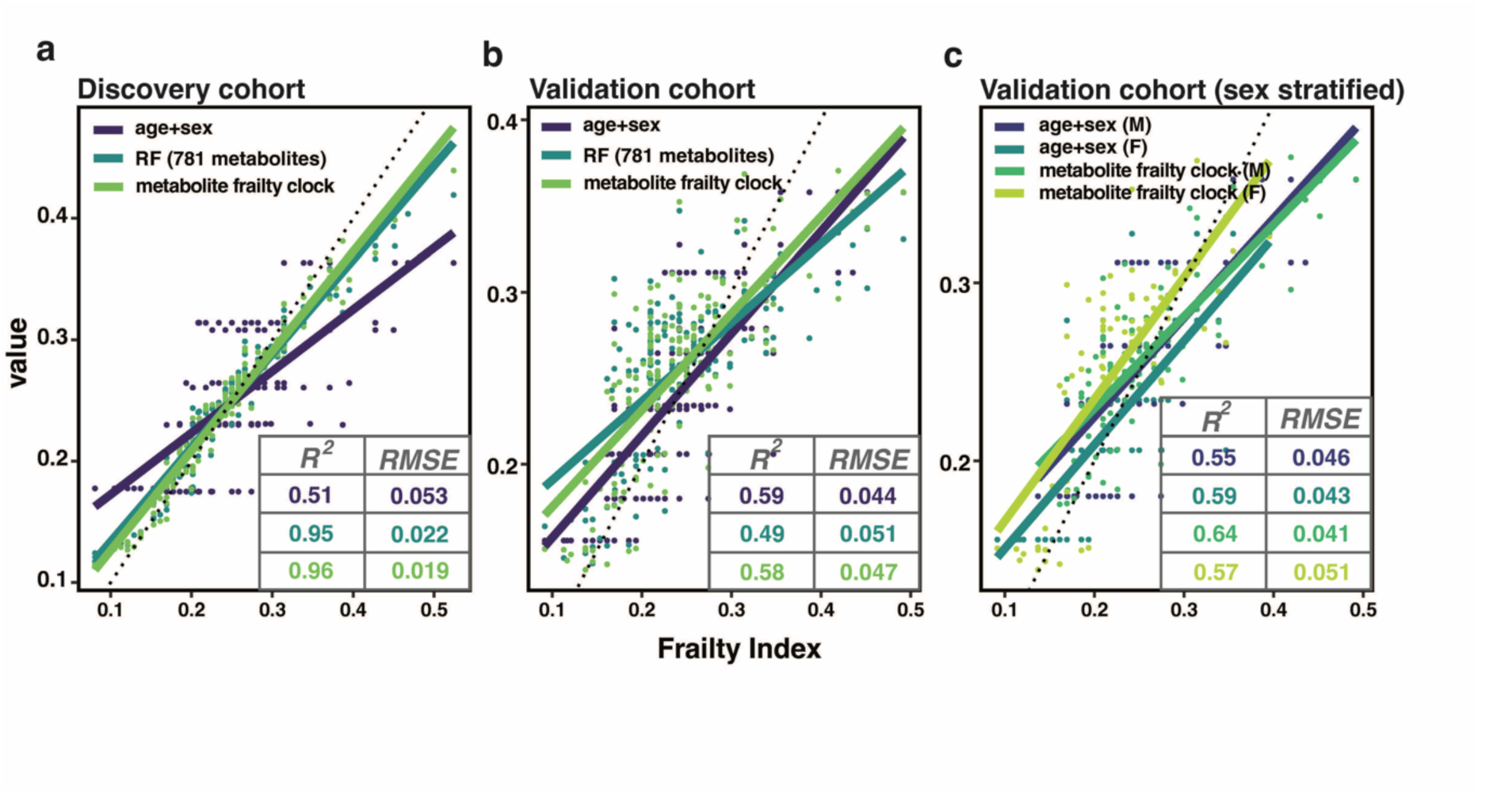
Performance of metabolite frailty clock in the discovery and validation cohorts. Frailty models were built via machine learning approaches, with frailty index scores as the dependent variable and three sets of variables as the independent variables: 1) The age+sex model, linear regression models using age and sex, trained in the discovery and validation cohorts respectively; 2) The RF (781 metabolites) model, a random forest model using all 781 metabolites detected in this study, age and sex; and 3) metabolite frailty clock model, a random forest model using 63 informativity-based metabolites, age and sex. The performance of models in the corresponding cohort/subcohort (Female, F and male, M) are presented using *R^2^*and Root-mean-square deviation (*RMSE*).

## Discussion

Using a longitudinal study of female and male mice, we identified both sex-independent and sex-specific metabolomic signatures of aging and frailty. Overall, we found that age related metabolites are enriched for lipid metabolism, while frailty related metabolites are enriched for amino acid metabolism and metabolism of cofactors and vitamins. B vitamin metabolism-related metabolites and lipid metabolism-related metabolites, respectively, are determined as candidate female- and male-specific frailty biomarkers.

### Age-related metabolites

Using a time course analysis in mice, we found a total of 527 metabolites significantly changed with age, representing the majority of measured metabolites. This result suggests dramatic change in the abundance of most metabolites in aging, and aligns with a previous longitudinal human study^21^. Among these age-related metabolites, we identified 86 hub metabolites by network analysis, and observed that these metabolites were enriched for lipid metabolism, including biosynthesis of unsaturated fatty acids, primary bile acid biosynthesis, and fatty acid elongation. Interestingly, several studies in mice have demonstrated manipulation of lipid metabolism as a method to extend longevity^35,36^. Our results using longitudinal data provide further evidence of the importance of lipid metabolism in aging. In terms of the individual metabolites changed in age, we identified 10 sphingomyelin species, which are of interest given the established link between sphingomyelins and longevity in humans^20^.

Additionally, we found that 42% of measured metabolites were significantly different between males and females, and the majority of these were also changed with age. Metabolites involved in aging and sex differences were enriched for a spectrum of pathways involved in lipid metabolism and digestive system. While sex differences in lipid metabolism have been widely recognised, our finding provides further evidence in the context of aging and aligns with the result from humans that the lipidome exhibits significant age-dependent differences between sexes^37^. Moreover, liver is the primary tissue for bile acid metabolism^38^, fatty acid metabolism^39^ and taurine metabolism (conjugation with bile acid^40^), all pathways identified in our study as displaying sex differences in aging. This suggests that the liver is strongly influenced by biological sex in aging, which aligns with transcriptomic results from our lab^41^. Furthermore, the presence of the mineral absorption and ferroptosis pathways in our findings^42,43^ reveal that the impact of aging on iron homeostasis^44^ is sex specific.

Sex stratified analysis revealed female specific metabolite markers of aging include amino acids and acylcarnitines. Previous work has extensively shown the role of amino acids and acylcarnitines in the regulation of aging^45,46^. Our results indicate significant changes of these metabolites in females in aging, but not necessarily in males, which is also implied in other studies^47^. As for the male specific aging biomarkers, we found phosphocholine (adjusted for sex) and spermine (only in males), both of which have been previously linked to overall aging^48,49^. Together, these findings give clues about how metabolic aging may occur differently in males and females.

### Frailty related metabolites

Although age-related metabolic markers are of interest, markers that are associated with health in aging may provide more clues about underlying mechanisms of the aging process, rather than the passing of time. To this end, we sought to identify metabolic features of frailty, a validated quantification of health in aging in both humans and mice. As frailty is strongly correlated with age, it was important that we identify metabolic markers of frailty, independent of age. We used a novel approach of calculating devFI, the deviation from the median frailty index of the corresponding age and sex group. In this way we are able to identify metabolites associated with individual variations in frailty at a given age and sex group, and distinguish these from metabolic changes that arise from aging. We applied a machine learning approach to select metabolites that are associated with both outcomes, FI and devFI. In the whole cohort study, we identified 149 metabolites features for devFI, among which 61 were not also associated with FI or age (**Fig.4b**). These include hippurate, a gut microbiome derived metabolite that has been previously associated with aging^50^. These metabolites are particularly interesting for further analysis as underlying markers of health, independent of age.

Overall, frailty-related metabolites were heavily enriched in amino acid metabolism and metabolism of cofactors and vitamins. The majority of the 20 proteinogenic amino acids metabolism pathways were enriched, suggesting amino acids serve as the main driver of frailty dynamics in mice. In humans, altered amino acid metabolism is also suggested to be associated with frailty^24^, in particular tryptophan metabolism^51^. Interestingly, nicotinate and nicotinamide metabolism were also over-represented in these candidate frailty biomarkers.

Recent work has shown that boosting nicotinamide levels is associated with improved health in aging, including improved frailty^34,52,53^. These results suggest pivotal differences in the metabolic mechanisms underlying aging and frailty.

Additionally, we applied linear mixed models to the metabolite features identified for frailty to look at their specific univariate association with frailty outcomes (**Fig.5a**). Notably, we found 23 metabolites that were associated with more than one frailty outcome (i.e., current FI, current devFI and/or change in FI over time) (**Fig.5d**), including ergothioneine, nicotinamide riboside (NR), phenyllactate, and creatinine. Interestingly, ergothionine is one of the most robust markers in our study, which is identified as FI, devFI and age-associated across both males and females. It has been previously identified as a frailty biomarker^54^, and is thought to promote healthy aging^55^. NR is an NAD precursor, part of the nicotinamide metabolism pathway, and boosting levels of NR are associated with improved health in aging^53,57^. Phenyllactate, is a catabolite of phenylalanine (phenylalanine metabolism is identified as enriched from frailty related metabolites) derived from *Lactobacillus*^58^, providing further evidence of the possible involvement of the microbiome in the development of frailty. Creatinine, a muscle breakdown product, has been associated with sarcopenia, functional limitation and frailty^59^. Additionally, alpha-ketoglutarate was the only metabolite for which the abundance in middle-aged mice was predictive of future frailty, suggesting it could be an early-biomarker of frailty, and/or a viable target for early intervention. In support of this, a recent study shows that alpha-ketoglutarate supplementation in mice reduced frailty^56^. Taken together, our results provide preclinical evidence for several potential biomarkers for frailty.

### Sex dimorphism in frailty

In order to identify sex-specific metabolic markers of frailty, we completed sex-stratified analysis. We identified vitamin B3/tryptophan metabolites, kynurenine and quinolinate, as being specifically associated with frailty in females. The findings for these two metabolites are consistent with previous studies^60,61^, where the link to frailty is sex-specific. FAD (vitamin B2), was also significantly associated with multiple frailty outcomes in females (**Supplementary Fig.9a**). FAD is one of the active forms of vitamin B2, however, previous studies in both sexes found that intake of vitamin B2 has no association with frailty^62,63^. Given our novel findings, we suggest further investigation into FAD as a female-specific marker of frailty. Another female-specific frailty biomarker is pyridoxate (vitamin B6), which is reported to be related to frailty^64^. In male mice, we identified mainly lipid metabolism-related metabolites, including sphingomyelins, three GPE species and one GPC species. These metabolites are lipid species that have been previously associated with frailty^65,66^ in both sexes, so the male specificity needs further investigation. Taken together, our results reveal evidence of sex specific biomarkers for frailty, and imply that B vitamin metabolism is a key feature of frailty development in females and lipid-related metabolism for males. We highly recommend applying the sex stratification approach in the future study of frailty biomarkers and mechanisms.

Importantly, and often ignored in other frailty biomarkers studies, we confirmed whether the same metabolites were associated with frailty outcomes in an independent validation cohort. Although the association of not all metabolites held in this cohort, we did find 9 metabolites showing persistent significance in the association with frailty outcomes, including ergothioneine and creatinine. Our validation cohort included mice that had long-term treatment with the NAD booster, NMN, suggesting that the association of these metabolites with frailty outcomes may be universal even under interventions, so these biomarkers should be investigated further.

Many ‘clocks’ have been built to predict chronological age based on either epigenetic or metabolomic features^67,68^. There is a growing focus, however, on building models to predict health- rather than age-related outcomes. Here, we build the first clock to directly predict frailty in mice. Our model performs extremely well in our discovery cohort, and although the performance of the frailty clock is similar to that of an age+sex only model (trained within the validation cohort) in females for the validation cohort, our clock outperforms the simple model in male samples. These results provide preliminary evidence that it is possible to predict frailty using metabolites, but suggest further work should be done in large datasets to develop a more universal metabolomics-based frailty clock.

There are some limitations to this study. Our validation dataset was relatively small, and mice were treated with NMN that may alter metabolite abundance levels. We suggest future work should validate these potential frailty biomarkers in larger cohorts, as well as in other mouse strains and humans. Additionally, the sample size for metabolomics data is relatively small, especially at the older ages, which might decrease the power of statistical analysis. Survival bias is also an issue to consider, as mice died over the course of the study and only those that were longest lived made it to timepoint 4 and 5. For future work, it will be ideal to conduct studies in a broader age range with an increased number of mice.

In summary, we performed the first longitudinal study of naturally aging female and male mice looking at metabolomics of frailty. We found aging related metabolites are mainly involved in lipid metabolism while frailty related metabolites are predominantly parts of amino acid metabolism and metabolism of cofactors and vitamins. Apart from whole cohort frailty biomarkers, we demonstrated the sex dimorphism in the associations between metabolite and frailty, and proposed sex specific frailty biomarkers.

## Material and methods

### Mice samples

Mice used in this study are from a larger intervention study, so detailed methods can be found in Kane et al (2024)^34^. Briefly, C57BL/6NIA mice, female (*n* = 40) and male (*n* = 47) were obtained from the National Institute on Aging (NIA) Aging Rodent Colony, among which, 20 female and 23 male mice were subjected to nicotinamide mononucleotide (NMN) treatment. Mice were group housed (4-5 mice per cage, although over the period of the experiment mice died and mice were left singly housed), at Harvard Medical School in ventilated microisolator cages, with a 12-hour light cycle, at 71°F with 45-50% humidity. Mice were fed AIN-93G Purified Rodent Diet (Dyets Inc, PA). All animal experiments were approved by the Institutional Animal Care and Use Committee of the Harvard Medical Area. In order to investigate aging and frailty related metabolites and mechanisms in naturally aging mice, we used non-NMN treated mice (female, *n* = 20; male, *n* = 24) as the discovery cohort for principal component analysis, feature selection, sex stratified analysis, association study, and metabolite frailty clock model building (**Fig.1**). We then tested the selected metabolite features and model in the NMN treated mice (validation cohort).

### Mouse Frailty assessment

Behavioral and clinical variables for clinical frailty index were measured in both the discovery and validation cohorts, at each time point (**Table1**). We utilized the mouse clinical frailty index^2^ (FI) that contains 31 health-related items for this study. Briefly, mice were scored either 0, 0.5 or 1 for the degree of deficit they showed in each item with 0 representing no deficit, 0.5 representing a mild deficit and 1 representing a severe deficit^69^. Apart from FI score itself, we introduced devFI score, that is the deviation of individual FI from the median FI for the corresponding sex, at the corresponding time point.

### Blood collection and processing

Mice were fasted for 5-6 hours, anesthetized with isoflurane (5%) and then blood was collected from the submandibular vein with a lancet (maximum 10% of mouse body weight, approx. 200- 300 ul), into a tube containing 20ul of 0.5M EDTA. Blood was mixed and stored on ice. Whole blood was centrifuged at 1500×g for 15 mins, plasma was removed and frozen at -80°C for subsequent metabolomics.

### Metabolites extraction, quantification and processing

Global metabolomics analysis was completed by Metabolon. Samples were prepared using the automated MicroLab STAR® system (Hamilton Company), and analyzed using Ultrahigh Performance Liquid Chromatography-Tandem Mass Spectroscopy (UPLC-MS/MS). We used raw metabolite data (peak area). We performed batch effects mitigation by calculating the mean metabolite value for the baseline time point across all mice, comparing it to the mean value of all other time points and excluding metabolites that presented the mean of the baseline significantly lower (0.05x, compared to the mean of all other time points) or significantly higher (10x compared to the mean of all other time points). We normalized each sample by dividing the metabolite value by the median of metabolite values for that sample to account for any collection batch effects and then derived the (natural) log-transformed values as the metabolite abundance. We derived 781 metabolites with greater than 5% unique abundance across samples.

### Metabolomics data variations

The number of metabolomics data is summarised in **Table1**. Metabolite abundance data was subjected to principal component (PC) analysis. We derived PC1 to PC10 and for each PC as the dependent variable, we applied linear regression models and obtained *p*-values, where we used mouse ID, time points, sex, cage (categorical variables), and age at assessment (continuous variables), respectively as the independent variable. Independent variables tested were then clustered according to Euclidean distance.

### Differential abundance analysis of metabolites

Log-transformed metabolite abundance data were subjected to differential abundance analysis by using the ‘limma’ pipeline with a spline. Briefly, metabolite abundance data was subjected to the limma time-course spline analysis, excluding time point 5 due to absence of female samples at this time point. We generated a matrix for a natural cubic spline based on the remaining time points, with degrees of freedom set at 3, and the matrix was used as the time factor. Design matrices for global differential abundance analysis included sex and sex by time interaction term, without assigning a reference level. The data along with the multi-factor design matrix were then subjected to linear modeling with the intra-block correlation based block on mouse ID, and empirical bayes smoothing of metabolite-wise standard deviations. We then determine metabolite abundance differences by defining a contrast matrix for each of the following four categories: 1) mixture of female and male samples, 2) female samples, 3) male samples, and 4) sex differences.

### Co-abundance analysis

We performed the co-abundance analysis of metabolites (excluding time point 5), with a soft threshold set at 9 to select metabolite abundance modules. For a given module, we derived the first principal component as the eigenvalue. To identify the association of metabolite modules with age, we applied a linear mixed model using the module eigenvalue as independent variable and age as dependent variable with adjustment for sex, allowing a random intercept for each mouse. P-values were then adjusted by the Bonferroni correction method. For each identified subset, metabolites that showed significance greater than 0.2 (correlation coefficient between the metabolite abundance and the age) and module membership greater than 0.8 were selected as the hub metabolites in the module.

### Pathway enrichment analysis

Metabolite set enrichment analysis was performed by using the hypergeometric test from R package *FELLA* (v. 1.20.0) to identify KEGG pathways that were overrepresented, with a cutoff of *p*-value set at 0.05.

### Feature selection

We applied a machine learning approach to identify FI/devFI related metabolites in the discovery cohort (**Fig. 4a**). We performed feature selection by fitting generalized linear regression models using the frailty assessment score (FI or devFI) as the dependent variable and the 781 metabolites abundance data as the independent variables, through a 100 x 5-fold cross validation approach. Briefly, we performed 100 runs of multivariate generalized regression with elastic net regularization. Within each run, the hyperparameters for the least Root mean square error (RMSE) were tuned using 5-fold cross-validation, and a list of metabolite features assigned a non-zero coefficient was derived. These lists (from 100 runs) were merged into a list of metabolite features, which were then ranked according to the importance, i.e. the presence percentage of the metabolites. We selected metabolites that made to the top 20% percentile as FI/devFI metabolites. FI is composed of the age-related base FI and devFI. Hence, we derived FI-age features by combining age metabolites (hub metabolites from co-abundance analysis) and FI metabolites, and devFI features by combining devFI metabolites with FI metabolites. We then obtained union features from the union of FI-age features and devFI features.

### Analysis of metabolite associated with frailty outcomes

For the association study, we applied mixed linear models allowing variations in individual mice as the random effect. We considered three outcomes for FI and devFI respectively (six in total), 1) the score at current age (age_c_), FI_c_ and devFI_c_; 2) a score at a future time point (age_f_), FI_f_ and devFI_f_; and 3) score change to a future time point, ᐃFI and ᐃdevFI (**Fig.5a**). We considered two scenarios in the analysis, where abundance of metabolites from a previous time point (age_p_) are: a) absent, only the current abundance of metabolites (MA_c_) is available. For each metabolite, we used one of the six score outcomes as the dependent variable and MA_c_ as the independent variable, adjusting for sex, age_c_ and age change from age_c_ to age_f_ (ᐃage_2_, only for outcomes 2) and 3)) in the non-interaction models. For interaction models, we considered abundance by age term and abundance by age change term; and b) present, the current and a previous abundance of metabolites and the age interval are available. We focused on metabolite abundance change (ᐃMA) in this scenario. We used one of the six score outcomes as the dependent variable and ᐃMA as the independent variable, adjusting for MA_c_, age change from age_p_ to age_c_ (ᐃage_1_), sex, age_c_ and age ᐃage_2_ (for outcomes 2) and 3)) in the non-interaction models. For interaction models, we included abundance change by age (and/or age change) terms and current abundance by age (and/or age change) term. devFI score is the deviation from the median at the age- and sex- specific group. Hence, current age was not included in the analyses of devFI in both non- and interaction models. The age variable used above was the actual days of assessment divided by 1,000, in order to be within the same scale.

### Metabolite frailty clock model building

After obtaining three sets of union features for the whole cohort, females, and males, we generated a single set of metabolite features from the above three sets. We ranked these features by occurrence frequency from the feature selection process (100 times repeated cross validation) within the whole cohort. Via a cross validation approach, we selected ‘mtry’ (the number of randomly drawn candidate variables out of which each split is selected when growing a tree) and the number of informativity-based top metabolite features that gave the least RMSE in predicting FI. The final metabolite frailty clock model was fit in the discovery cohort, constructed using a random forest regression with FI as the dependent variable and the top metabolites features, age and sex as the independent variables. We also fit linear regression models with age and sex as the independent variables in respective the discovery and validation cohort, and a random forest model using all the 781 metabolites, age and sex in the discovery cohort for comparative purposes.

### Statistics

All statistical analyses were performed using R (version 4.3.0). Differentially abundant metabolites (DAMs) are selected by controlling for a 5% Benjamini-Hochberg false discovery rate (adjusted *p*-values < 0.05). For univariate association study, the significance was determined by controlling for a 5% Benjamini-Hochberg false discovery rate (adjusted *p*-values < 0.05).

## Data availability

Mice metadata, metabolite abundance data, and R markdown file for data analysis are available at https://github.com/Kane-Lab-ISB/longitudinal-metabolite-analysis-in-mice.git.

## Author contributions

A.E.K. and D.A.S. conceived and designed the study. A.E.K. performed the experiments. D.Z. conducted the data analysis, with contribution from J.Z.W., P.G., and B.A.S. P.G. provided critical feedback. D.Z., J.Z.W., and A.E.K. drafted and revised the manuscript with help from all authors. All authors have read and agreed to the published version of the manuscript.

## Correspondence

Correspondence to Alice E. Kane.

## References

1. Mitnitski, A., Howlett, S. E. & Rockwood, K. Heterogeneity of Human Aging and Its Assessment. J. Gerontol. A Biol. Sci. Med. Sci. 72, 877–884 (2017).

2. Whitehead, J. C. et al. A clinical frailty index in aging mice: comparisons with frailty index data in humans. J. Gerontol. A Biol. Sci. Med. Sci. 69, 621–632 (2014).

3. Graber, T. G., Ferguson-Stegall, L., Liu, H. & Thompson, L. V. Voluntary Aerobic Exercise Reverses Frailty in Old Mice. J. Gerontol. A Biol. Sci. Med. Sci. 70, 1045–1058 (2015).

4. Mitnitski, A. B., Mogilner, A. J. & Rockwood, K. Accumulation of deficits as a proxy measure of aging. ScientificWorldJournal 1, 323–336 (2001).

5. Heinze-Milne, S. D., Banga, S. & Howlett, S. E. Frailty Assessment in Animal Models. Gerontology 65, 610–619 (2019).

6. Bisset, E. S. & Howlett, S. E. The biology of frailty in humans and animals: Understanding frailty and promoting translation. Aging Med (Milton) 2, 27–34 (2019).

7. Kane, A. E. & Sinclair, D. A. Frailty biomarkers in humans and rodents: Current approaches and future advances. Mech. Ageing Dev. 180, 117–128 (2019).

8. Pan, Y., Ji, T., Li, Y. & Ma, L. Omics biomarkers for frailty in older adults. Clin. Chim. Acta. 510, 363–372 (2020).

9. López-Otín, C., Blasco, M. A., Partridge, L., Serrano, M. & Kroemer, G. Hallmarks of aging: An expanding universe. Cell 186, 243–278 (2023).

10. Kalyani, R. R., Varadhan, R., Weiss, C. O., Fried, L. P. & Cappola, A. R. Frailty status and altered glucose-insulin dynamics. J. Gerontol. A Biol. Sci. Med. Sci. 67, 1300–1306 (2012).

11. Sinclair, A. J. & Abdelhafiz, A. H. Metabolic Impact of Frailty Changes Diabetes Trajectory. Metabolites 13, (2023).

12. Panyard, D. J., Yu, B. & Snyder, M. P. The metabolomics of human aging: Advances, challenges, and opportunities. Sci Adv 8, eadd6155 (2022).

13. Mutlu, A. S., Duffy, J. & Wang, M. C. Lipid metabolism and lipid signals in aging and longevity. Dev. Cell 56, 1394–1407 (2021).

14. Dunn, W. B. et al. Molecular phenotyping of a UK population: defining the human serum metabolome. Metabolomics 11, 9–26 (2015).

15. Rist, M. J. et al. Metabolite patterns predicting sex and age in participants of the Karlsruhe Metabolomics and Nutrition (KarMeN) study. PLoS One 12, e0183228 (2017).

16. Dato, S. et al. Amino acids and amino acid sensing: implication for aging and diseases. Biogerontology 20, 17–31 (2019).

17. Weichhart, T. mTOR as Regulator of Lifespan, Aging, and Cellular Senescence: A Mini- Review. Gerontology 64, 127–134 (2018).

18. Yu, Z. et al. Human serum metabolic profiles are age dependent. Aging Cell 11, 960–967 (2012).

19. Peters, K. et al. Metabolic drift in the aging nervous system is reflected in human cerebrospinal fluid. Sci. Rep. 11, 18822 (2021).

20. Mielke, M. M. et al. Factors affecting longitudinal trajectories of plasma sphingomyelins: the Baltimore Longitudinal Study of Aging. Aging Cell 14, 112–121 (2015).

21. Darst, B. F., Koscik, R. L., Hogan, K. J., Johnson, S. C. & Engelman, C. D. Longitudinal plasma metabolomics of aging and sex. Aging 11, 1262–1282 (2019).

22. Mishra, M., Wu, J., Kane, A. E. & Howlett, S. E. The intersection of frailty and metabolism. Cell Metab. 36, 893–911 (2024).

23. Cesari, M., Calvani, R. & Marzetti, E. Frailty in Older Persons. Clin. Geriatr. Med. 33, 293– 303 (2017).

24. Calvani, R. et al. Amino Acid Profiles in Older Adults with Frailty: Secondary Analysis from MetaboFrail and BIOSPHERE Studies. Metabolites 13, (2023).

25. Gordon, E. H. et al. Sex differences in frailty: A systematic review and meta-analysis. Exp. Gerontol. 89, 30–40 (2017).

26. Yu, H., Armstrong, N., Pavela, G. & Kaiser, K. Sex and Race Differences in Obesity- Related Genetic Susceptibility and Risk of Cardiometabolic Disease in Older US Adults. JAMA Netw Open 6, e2347171 (2023).

27. de Ritter, R. et al. Sex differences in the risk of vascular disease associated with diabetes. Biol. Sex Differ. 11, 1 (2020).

28. Bell, J. A. et al. Sex differences in systemic metabolites at four life stages: cohort study with repeated metabolomics. BMC Med. 19, 58 (2021).

29. Zhao, L., Mao, Z., Woody, S. K. & Brinton, R. D. Sex differences in metabolic aging of the brain: insights into female susceptibility to Alzheimer’s disease. Neurobiol. Aging 42, 69–79 (2016).

30. Varghese, M., Song, J. & Singer, K. Age and Sex: Impact on adipose tissue metabolism and inflammation. Mech. Ageing Dev. 199, 111563 (2021).

31. Costanzo, M. et al. Sex differences in the human metabolome. Biol. Sex Differ. 13, 30 (2022).

32. Hägg, S. & Jylhävä, J. Sex differences in biological aging with a focus on human studies. Elife 10, (2021).

33. Hägg, S., Jylhävä, J., Wang, Y., Czene, K. & Grassmann, F. Deciphering the genetic and epidemiological landscape of mitochondrial DNA abundance. Hum. Genet. 140, 849–861 (2021).

34. 34. Kane, A. E., et al. Long-term NMN treatment increases lifespan and healthspan in mice in a sex dependent manner. *bioRxiv* (2024) doi:10.1101/2024.06.21.599604.

35. Canaan, A. et al. Extended lifespan and reduced adiposity in mice lacking the FAT10 gene. Proc. Natl. Acad. Sci. U. S. A. 111, 5313–5318 (2014).

36. Streeper, R. S. et al. Deficiency of the lipid synthesis enzyme, DGAT1, extends longevity in mice. Aging 4, 13–27 (2012).

37. Tabassum, R. et al. Lipidome- and Genome-Wide Study to Understand Sex Differences in Circulatory Lipids. J. Am. Heart Assoc. 11, e027103 (2022).

38. Phelps, T., Snyder, E., Rodriguez, E., Child, H. & Harvey, P. The influence of biological sex and sex hormones on bile acid synthesis and cholesterol homeostasis. Biol. Sex Differ. 10, 52 (2019).

39. Palmisano, B. T., Zhu, L., Eckel, R. H. & Stafford, J. M. Sex differences in lipid and lipoprotein metabolism. Mol Metab 15, 45–55 (2018).

40. Sjovall, J. Dietary glycine and taurine on bile acid conjugation in man; bile acids and steroids 75. Proc. Soc. Exp. Biol. Med. 100, 676–678 (1959).

41. Zhu, D. et al. Sex dimorphism and tissue specificity of gene expression changes in aging mice. Biol. Sex Differ. 15, 89 (2024).

42. Dixon, S. J. et al. Ferroptosis: an iron-dependent form of nonapoptotic cell death. Cell 149, 1060–1072 (2012).

43. Lindeman, R. D. Mineral metabolism in the aging and the aged. J. Am. Coll. Nutr. 1, 49–73 (1982).

44. Zeidan, R. S., Han, S. M., Leeuwenburgh, C. & Xiao, R. Iron homeostasis and organismal aging. Ageing Res. Rev. 72, 101510 (2021).

45. Austad, S. N., Smith, J. R. & Hoffman, J. M. Amino acid restriction, aging, and longevity: an update. Front Aging 5, 1393216 (2024).

46. Jarrell, Z. R. et al. Plasma acylcarnitine levels increase with healthy aging. Aging 12, 13555–13570 (2020).

47. Sol, J. et al. Plasma acylcarnitines and gut-derived aromatic amino acids as sex-specific hub metabolites of the human aging metabolome. Aging Cell 22, e13821 (2023).

48. Jové, M. et al. Human Aging Is a Metabolome-related Matter of Gender. J. Gerontol. A Biol. Sci. Med. Sci. 71, 578–585 (2016).

49. Xu, T.-T. et al. Spermidine and spermine delay brain aging by inducing autophagy in SAMP8 mice. Aging 12, 6401–6414 (2020).

50. De Simone, G., Balducci, C., Forloni, G., Pastorelli, R. & Brunelli, L. Hippuric acid: Could became a barometer for frailty and geriatric syndromes? Ageing Res Rev 72, 101466 (2021).

51. Al Saedi, A., Chow, S., Vogrin, S., Guillemin, G. J. & Duque, G. Association Between Tryptophan Metabolites, Physical Performance, and Frailty in Older Persons. Int. J. Tryptophan Res. 15, 11786469211069951 (2022).

52. Zhang, H. et al. NAD^+^ repletion improves mitochondrial and stem cell function and enhances life span in mice. Science 352, 1436–1443 (2016).

53. Mills, K. F. et al. Long-Term Administration of Nicotinamide Mononucleotide Mitigates Age- Associated Physiological Decline in Mice. Cell Metab. 24, 795–806 (2016).

54. Kameda, M., Teruya, T., Yanagida, M. & Kondoh, H. Frailty markers comprise blood metabolites involved in antioxidation, cognition, and mobility. Proc. Natl. Acad. Sci. U. S. A. 117, 9483–9489 (2020).

55. Katsube, M. et al. Ergothioneine promotes longevity and healthy aging in male mice. Geroscience (2024) doi:10.1007/s11357-024-01111-5.

56. Asadi Shahmirzadi, A., et al. Alpha-Ketoglutarate, an Endogenous Metabolite, Extends Lifespan and Compresses Morbidity in Aging Mice. Cell Metab. 32, 447–456.e6 (2020).

57. Imai, S.-I. & Guarente, L. NAD+ and sirtuins in aging and disease. Trends Cell Biol 24, 464–471 (2014).

58. Preidis, G. A. et al. Microbial-Derived Metabolites Reflect an Altered Intestinal Microbiota during Catch-Up Growth in Undernourished Neonatal Mice. J. Nutr. 146, 940–948 (2016).

59. Shlipak, M. G. et al. The presence of frailty in elderly persons with chronic renal insufficiency. Am J Kidney Dis 43, 861–867 (2004).

60. 60. Westbrook, R., et al. Kynurenines link chronic inflammation to functional decline and physical frailty. JCI Insight 5, (2020).

61. Chung, T. et al. Deletion of quinolinate phosphoribosyltransferase gene accelerates frailty phenotypes and neuromuscular decline with aging in a sex-specific pattern. Aging Cell 22, e13849 (2023).

62. Cheng, X. et al. Association between B-vitamins intake and frailty among patients with chronic obstructive pulmonary disease. Aging Clin. Exp. Res. 35, 793–801 (2023).

63. Balboa-Castillo, T. et al. Low vitamin intake is associated with risk of frailty in older adults. Age Ageing 47, 872–879 (2018).

64. Kato, N. et al. Relationship of Low Vitamin B6 Status with Sarcopenia, Frailty, and Mortality: A Narrative Review. Nutrients 16, (2024).

65. Laurila, P.-P. et al. Sphingolipids accumulate in aged muscle, and their reduction counteracts sarcopenia. Nat Aging 2, 1159–1175 (2022).

66. Marron, M. M., Yao, S., Shah, R. V., Murthy, V. L. & Newman, A. B. Metabolomic characterization of vigor to frailty among community-dwelling older Black and White men and women. Geroscience 46, 2371–2389 (2024).

67. Mutz, J., Iniesta, R. & Lewis, C. M. Metabolomic age (MileAge) predicts health and life span: A comparison of multiple machine learning algorithms. Sci Adv 10, eadp3743 (2024).

68. Mak, J. K. L. et al. Temporal Dynamics of Epigenetic Aging and Frailty From Midlife to Old Age. J Gerontol A Biol Sci Med Sci 79, (2024).

69. Kane, A. E. et al. A Comparison of Two Mouse Frailty Assessment Tools. J. Gerontol. A Biol. Sci. Med. Sci. 72, 904–909 (2017).

