## Supplemental Figures for "Metabolomics biomarkers of frailty: a longitudinal study of aging female and male mice"

Title:

**Supplementary Figures**

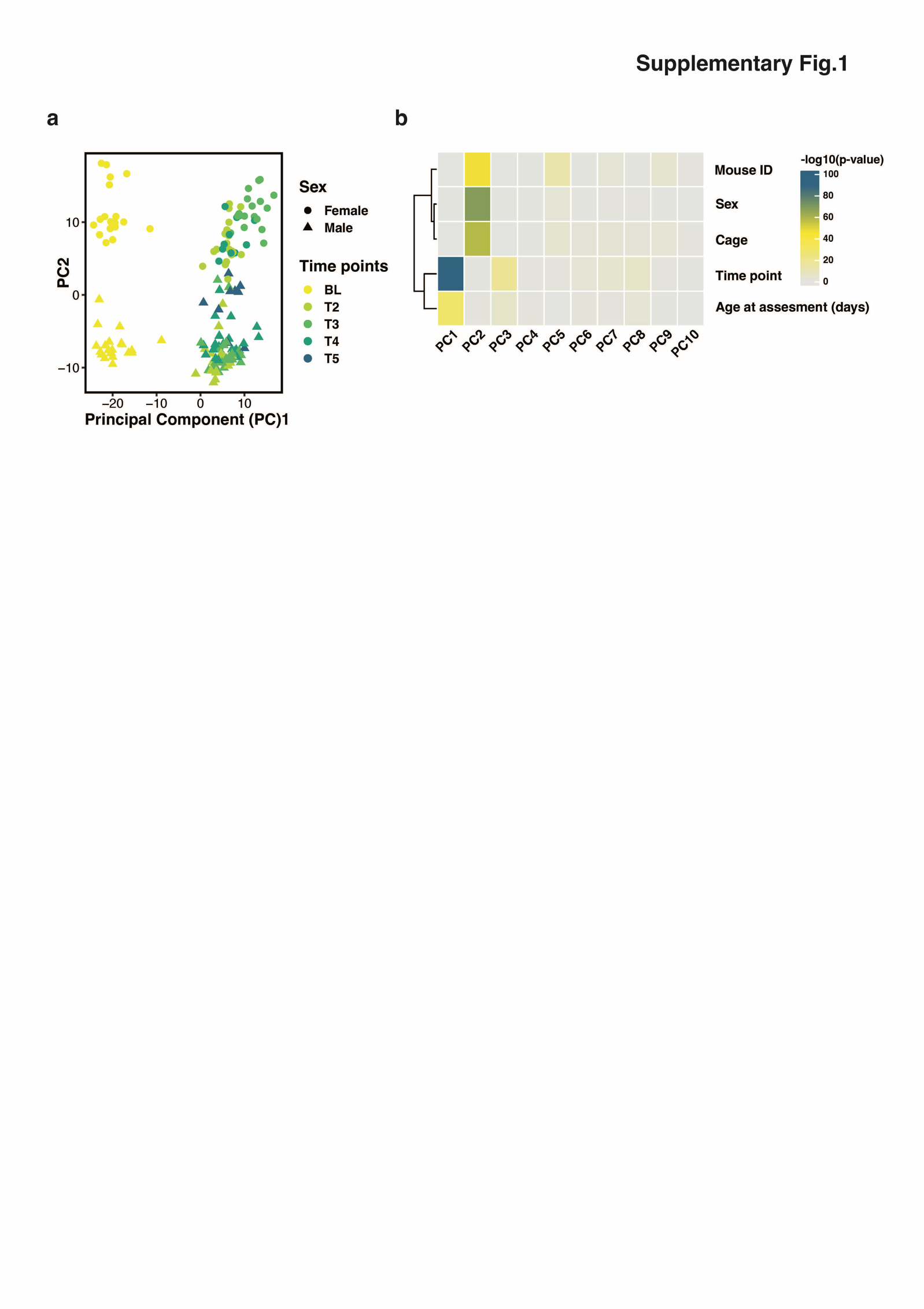

### Supplementary Fig.1. Principal component (PC) analysis of metabolic data.

**(a)** Scatter plot of PC1 and PC2 stratified by 5 time points (color) and sexes (shape). **(b)** The respective association of PC1 to PC10 with selected factors by linear regressions. The factors are hierarchically clustered, and the significance levels are presented by the color scale.

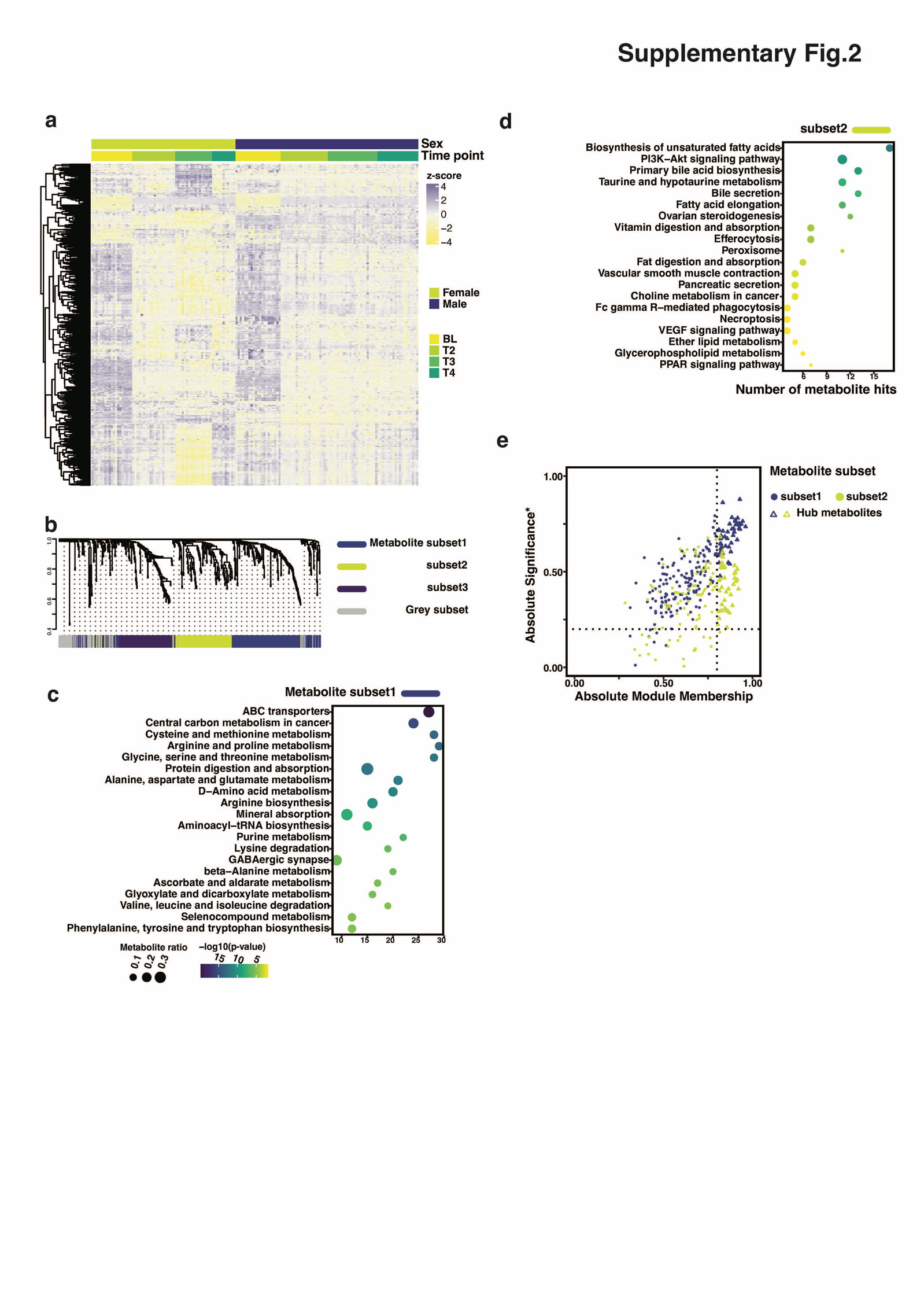

### Supplementary Fig.2. Analysis of sex independent age-related differentially abundant metabolites.

Differential abundance analysis was performed using all samples (excluding time point T5) in the study. 527 sex-independent age-related differentially abundant metabolites DAMs were selected from comparisons of the mixture of female and male samples (sex independent) at different time points and by controlling for a 5% Benjamini-Hochberg false discovery rate (adjusted *p*-values < 0.05). **(a)** Heatmap of z-scores of metabolite abundance for both sexes at four time points. **(b)** Hierarchical clustering dendrogram of co-abundant metabolites in modules. Over-represented pathways from the metabolites within subset1 **(c)** and subset2 **(d)** that presented significant change over the time course. The number of hits (metabolite) from the hub metabolites set is shown by x-axis, ratio of the hit number to total metabolites in the enriched pathway is represented by dot size and p-value is colored by levels. **(e)** Selection of hub metabolites in the two subsets based on significance (correlation of metabolite abundance with age) and module membership.

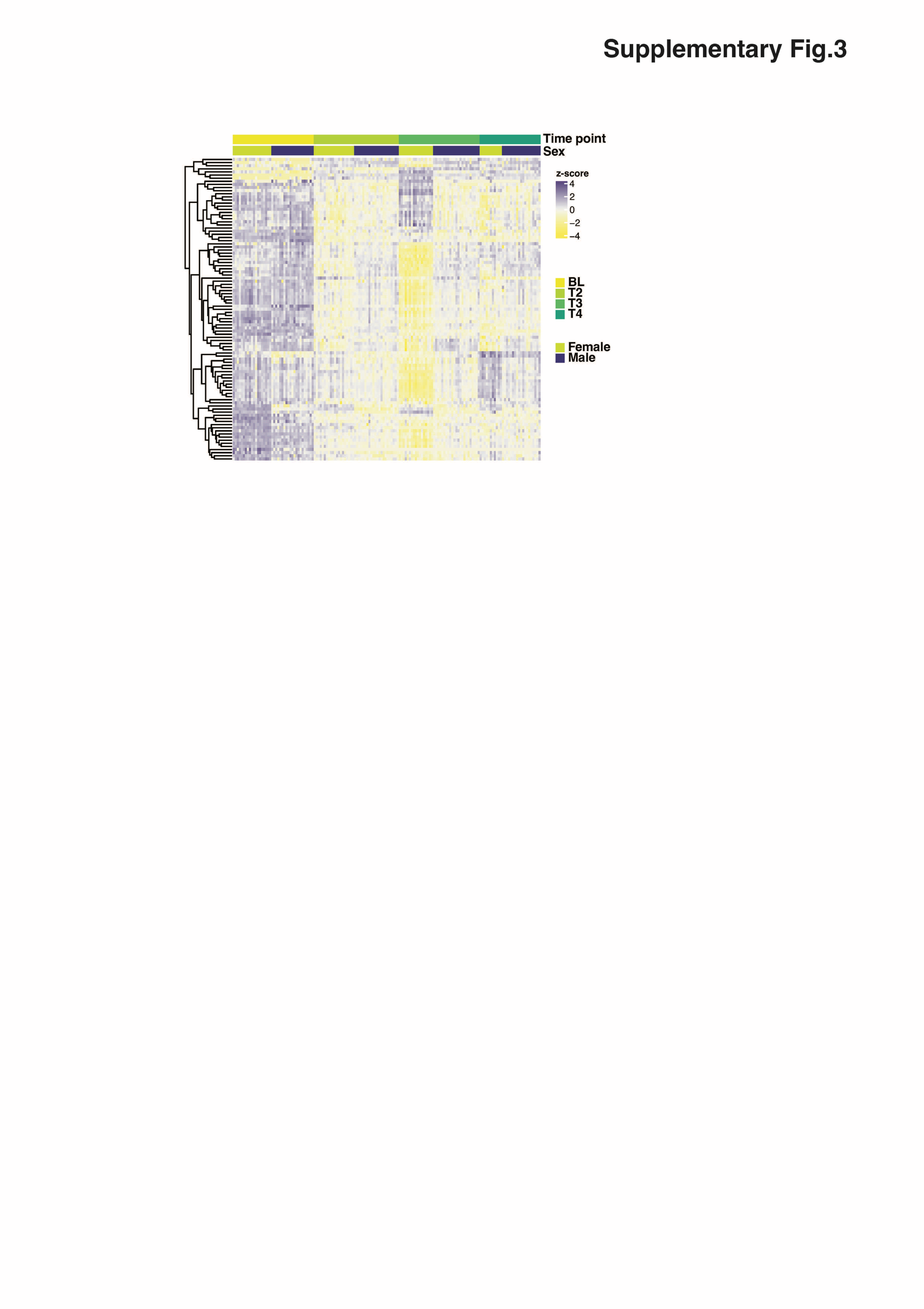

### Supplementary Fig.3. Metabolite abundance for 97 common age-related metabolites from four groups.

Differential abundance analysis was performed using all samples (excluding time point T5) in the study. Age-related differentially abundant metabolites (DAMs) were determined by comparisons within four groups: the mixture of females and males (sex independent), female specific, male specific, and sex differences, and by controlling for a 5% Benjamini-Hochberg false discovery rate (adjusted *p*-values < 0.05). Heatmap of metabolite abundance for 97 common DAMs from the four groups including both sexes at four time points were shown.

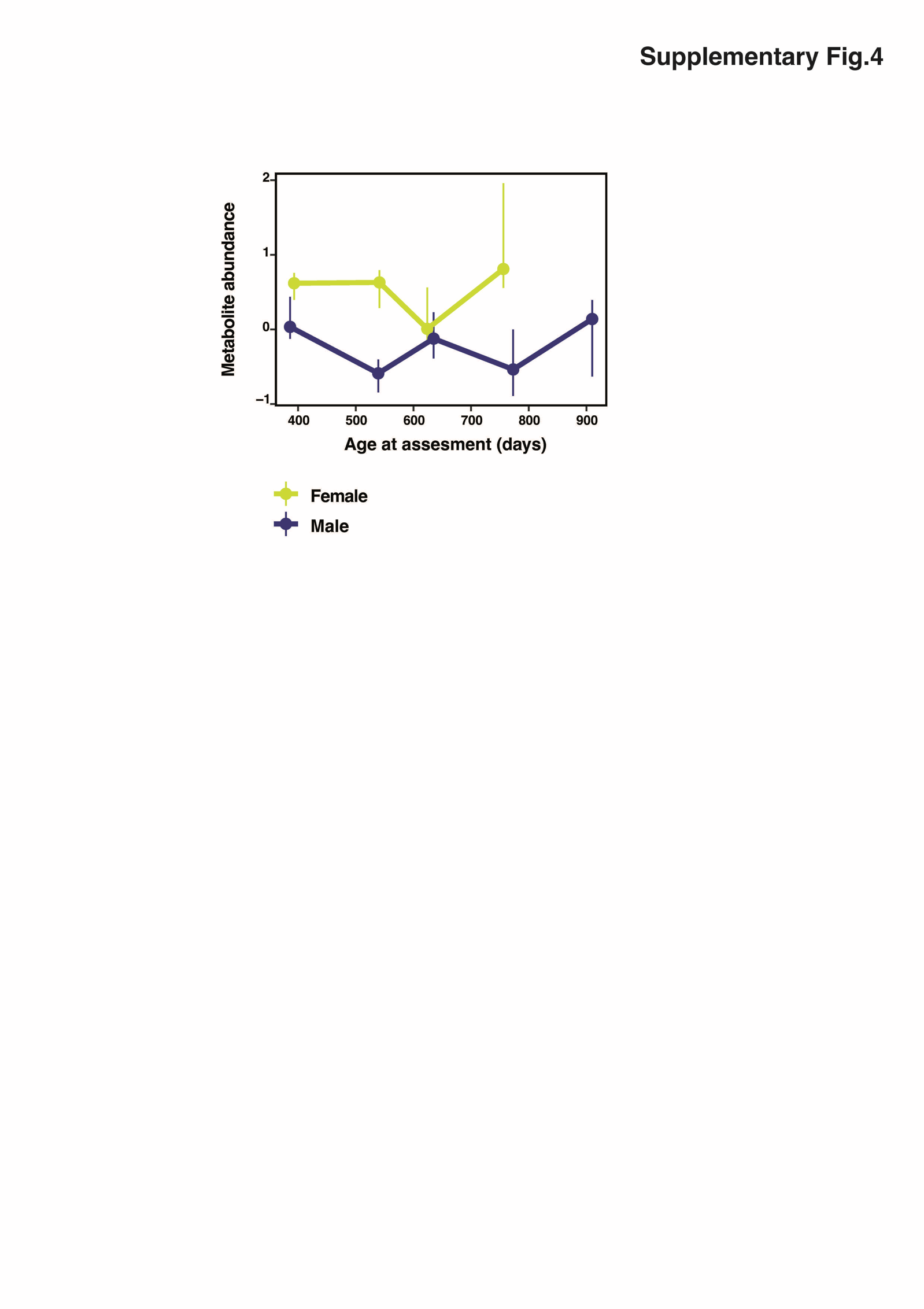

### Supplementary Fig.4. Abundance of corticosterone in the longitudinal study.

Abundance of corticosterone in females and males at time points. Error bars with dots show bottom quartile, median and top quartile of the values at each sex and age group.

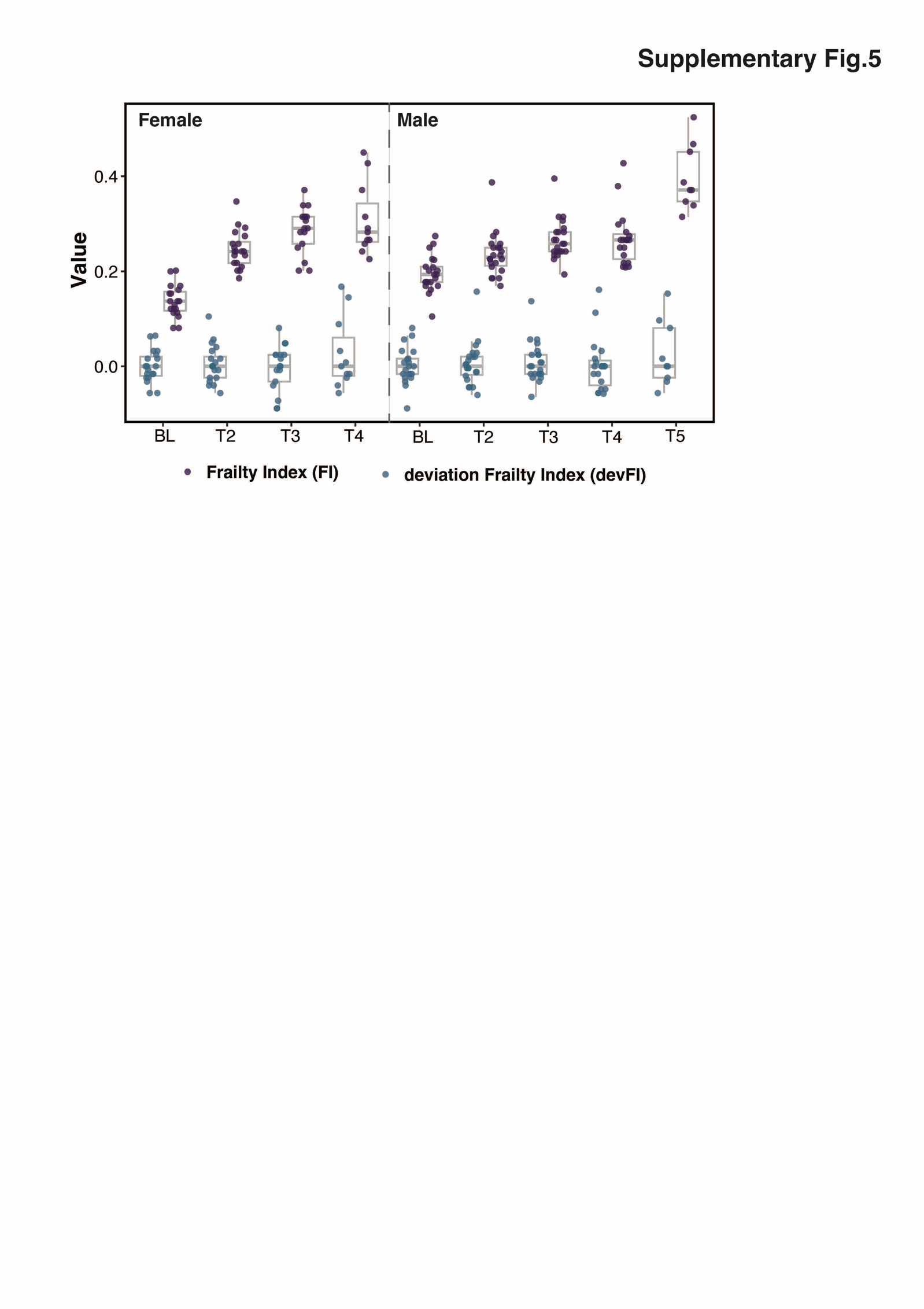

### Supplementary Fig.5. Frailty index and deviation frailty index scores in the discovery cohort.

Boxplots and jitter plots showing frailty index and deviation frailty index at five time points for females and males in the discovery cohort

.

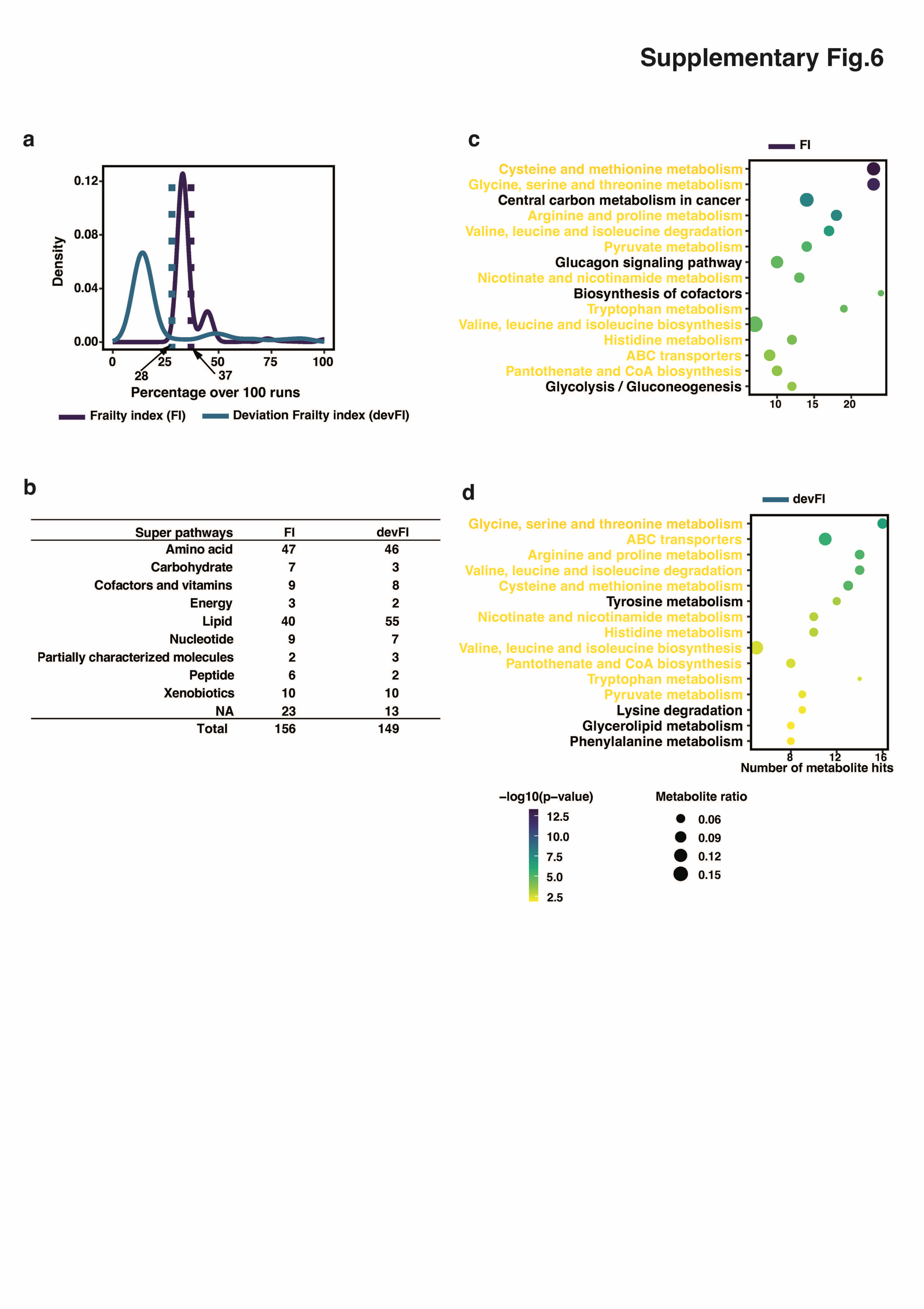

### Supplementary Fig.6. Selection of metabolites for frailty index and deviation frailty index.

Feature selection was performed using Elastic net regularization via a 100 times repeated 5-fold cross validation approach. **(a)** Density plot showing the distribution of presence ratio and the cutoff threshold for the selection. **(b)** List of number of FI/devFI metabolites in super pathways of metabolites. Over-represented pathways by metabolic data enrichment analysis for frailty index **(c)** and deviation frailty index **(d)**. Over-represented pathways from the metabolites within subset1 **(c)** and subset2 **(d)** that presented significant change over the time course. The number of hits (metabolite) from the hub metabolites set is shown by x-axis, ratio of the hit number to total metabolites in the enriched pathway is represented by dot size and p-value is colored by levels.

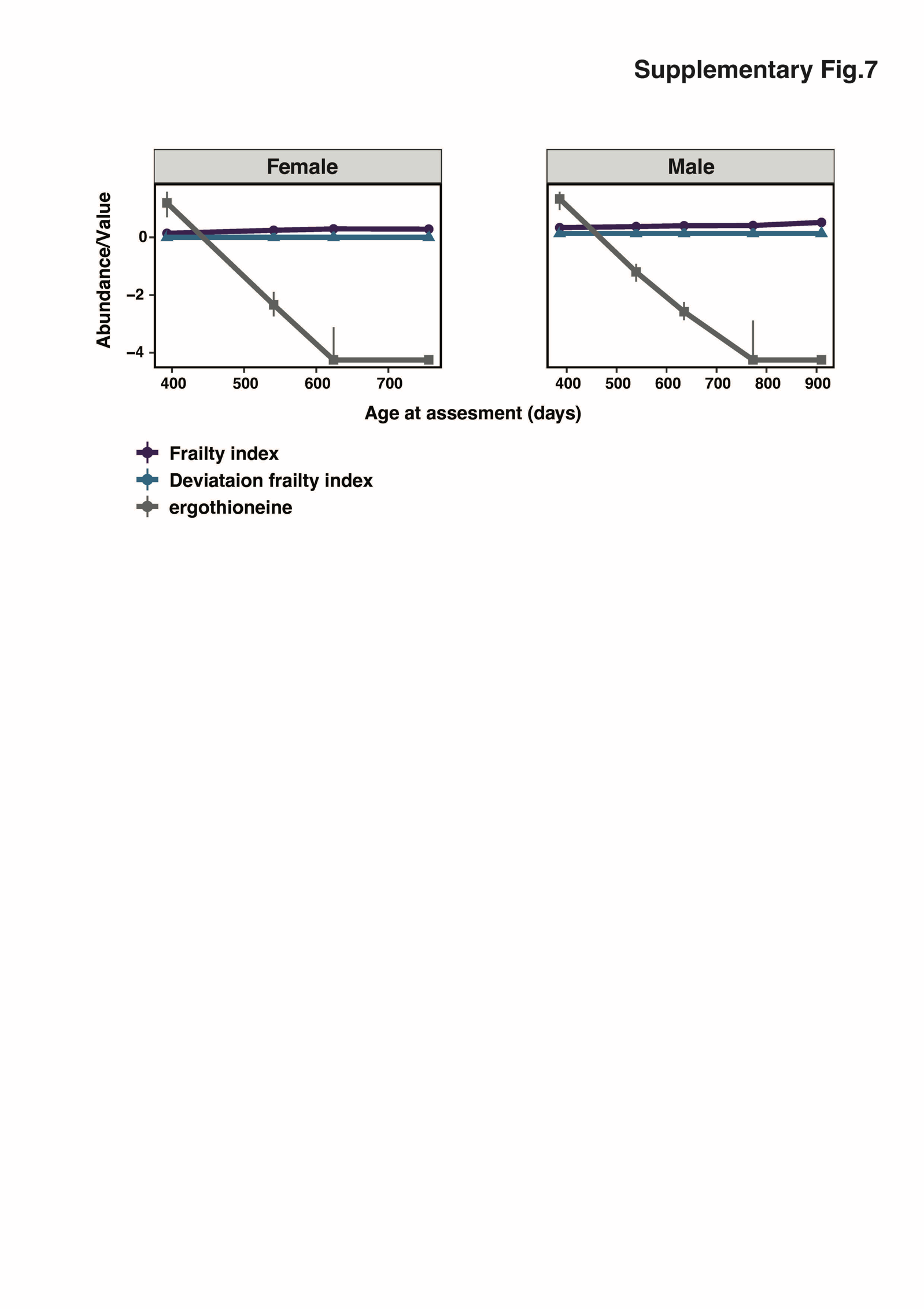

### Supplementary Fig.7. Dynamics of frailty index, deviation frailty index and ergothioneine over the time course.

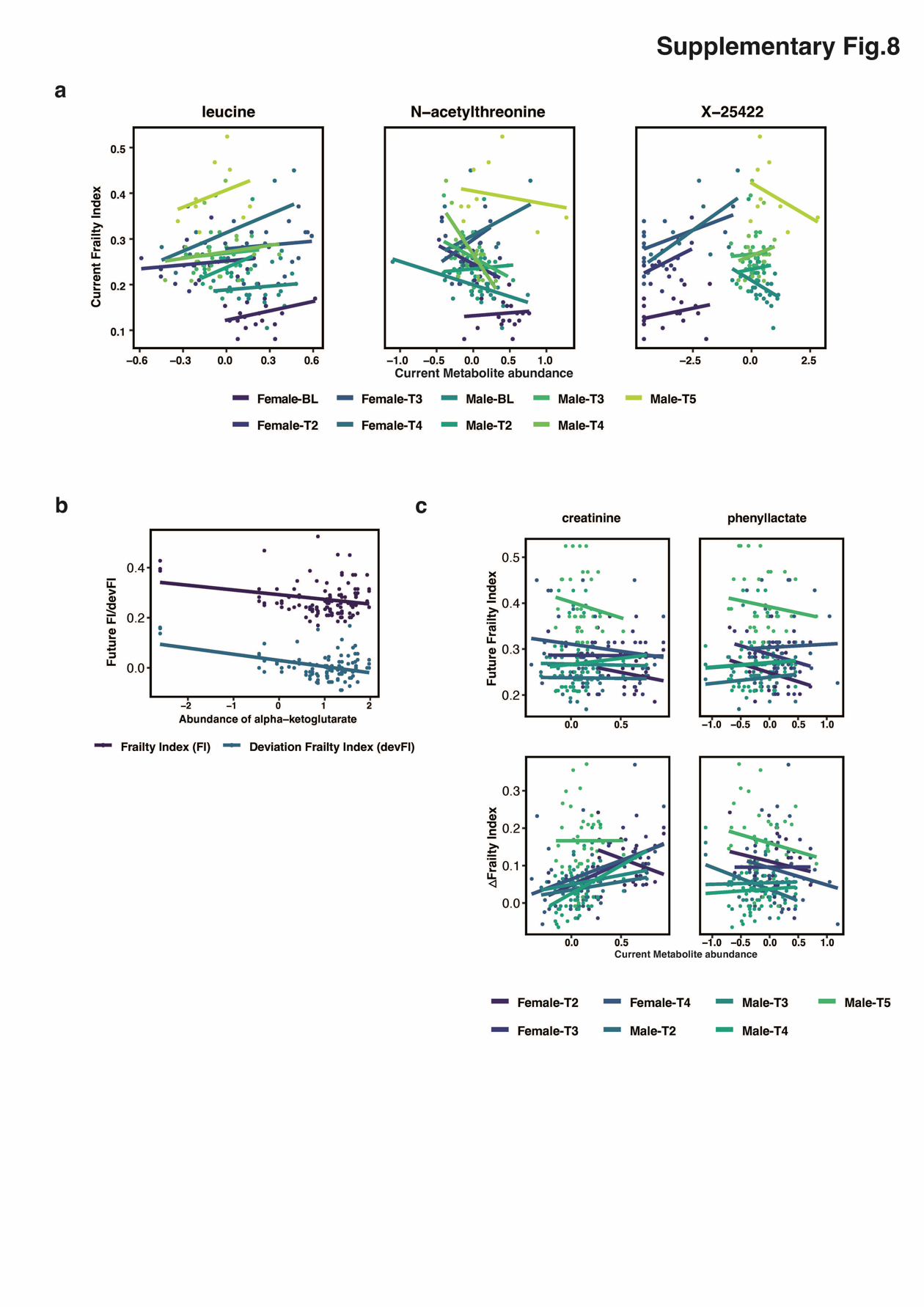

### Supplementary Fig.8. Associations between metabolites with frailty outcomes in different age- and sex- specific groups.

**(a)** Scatter plots showing associations of leucine, N-acetylthreonine and X-25422 with current frailty index at 5 time points in female and males. **(b)** Scatter plot presenting the relationships between abundance of alpha-ketoglutarate at baseline time point and future frailty index (FI) and future deviation frailty index (devFI). **(c)** Scatter plots showing relationships between the abundance of at current age and future FI and FI changes (ᐃFrailty Index). While there are no clear relationships between current metabolite abundance and future FI, creatinine and phenyllactate present associations with FI change.

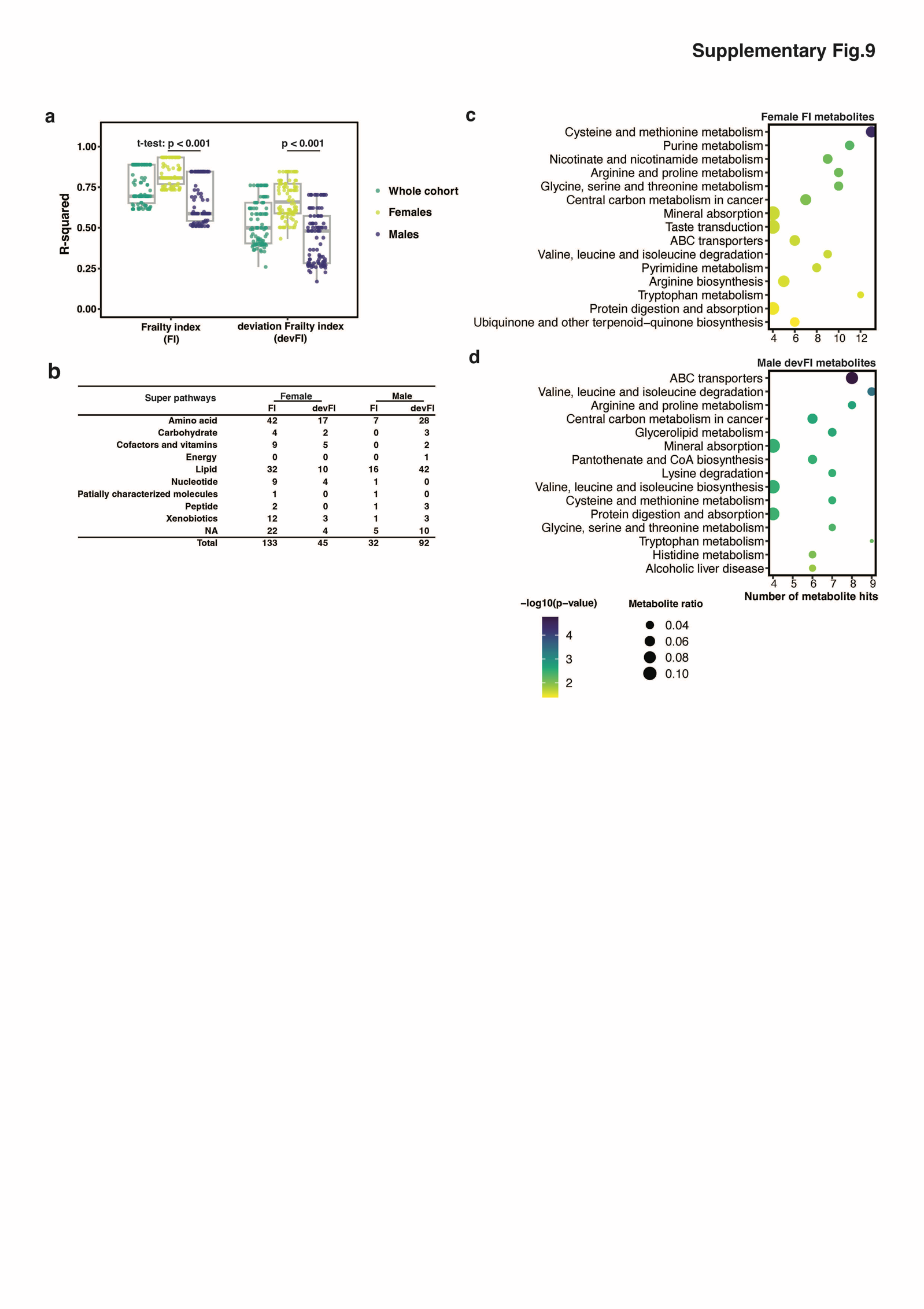

### Supplementary Fig.9. Selection of sex specific metabolites for frailty index and deviation frailty index.

During the feature selection procedure, for each run, performance of the model (root-mean-square error) in the whole cohort, female samples and male samples was derived. **(a)** boxplot showing the differences in performance. **(b)** List of number of FI/devFI metabolites in super pathways of metabolites in females and males respectively. Over-represented pathways by metabolic data enrichment analysis for female specific FI metabolites **(c)** and male specific devFI metabolites **(d)**. The number of hits (metabolite) from the hub metabolites set is shown by x-axis, ratio of the hit number to total metabolites in the enriched pathway is represented by dot size and p-value is colored by levels.

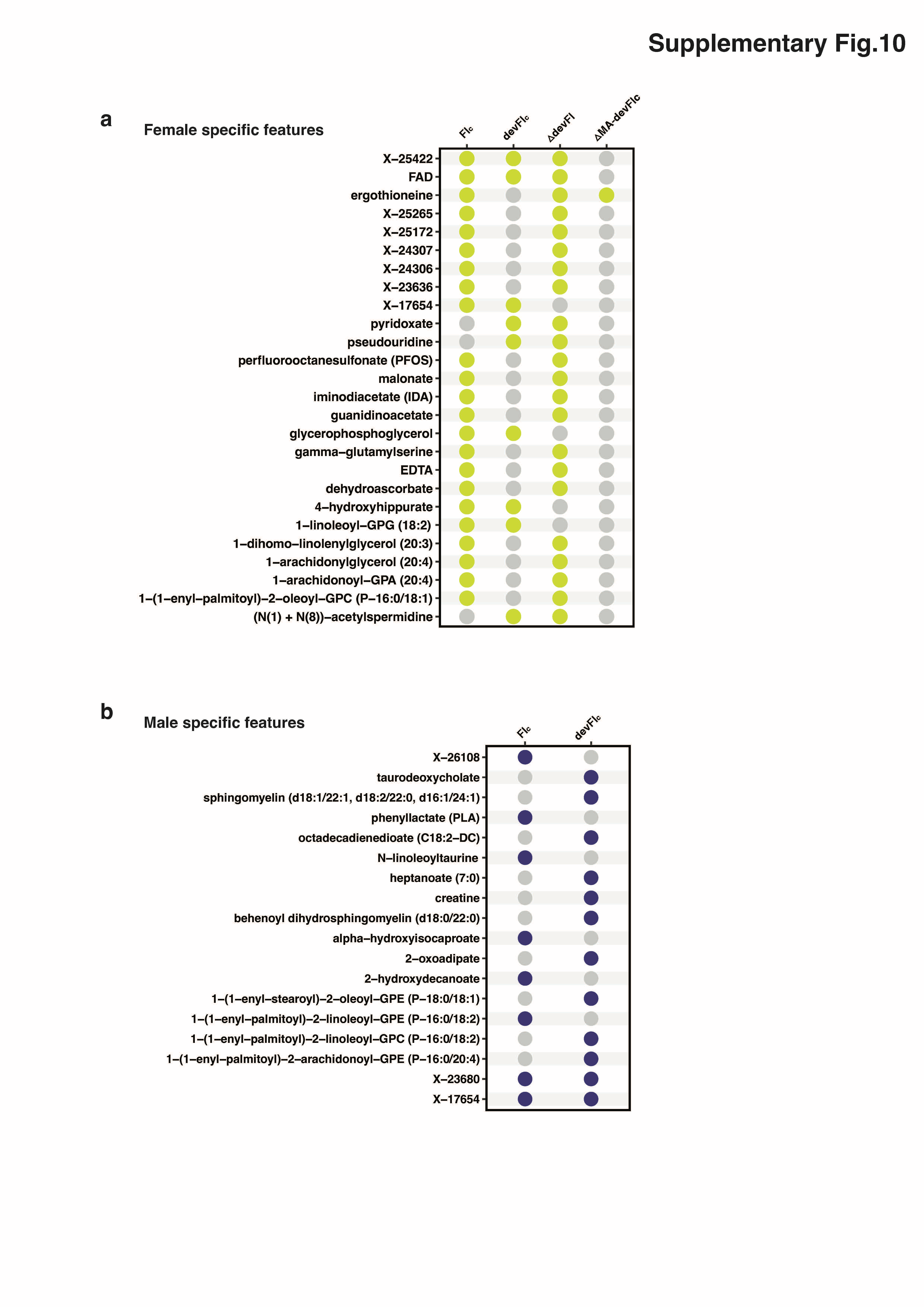

### Supplementary Fig.10. Common metabolites that present significance in the sex specific association study.

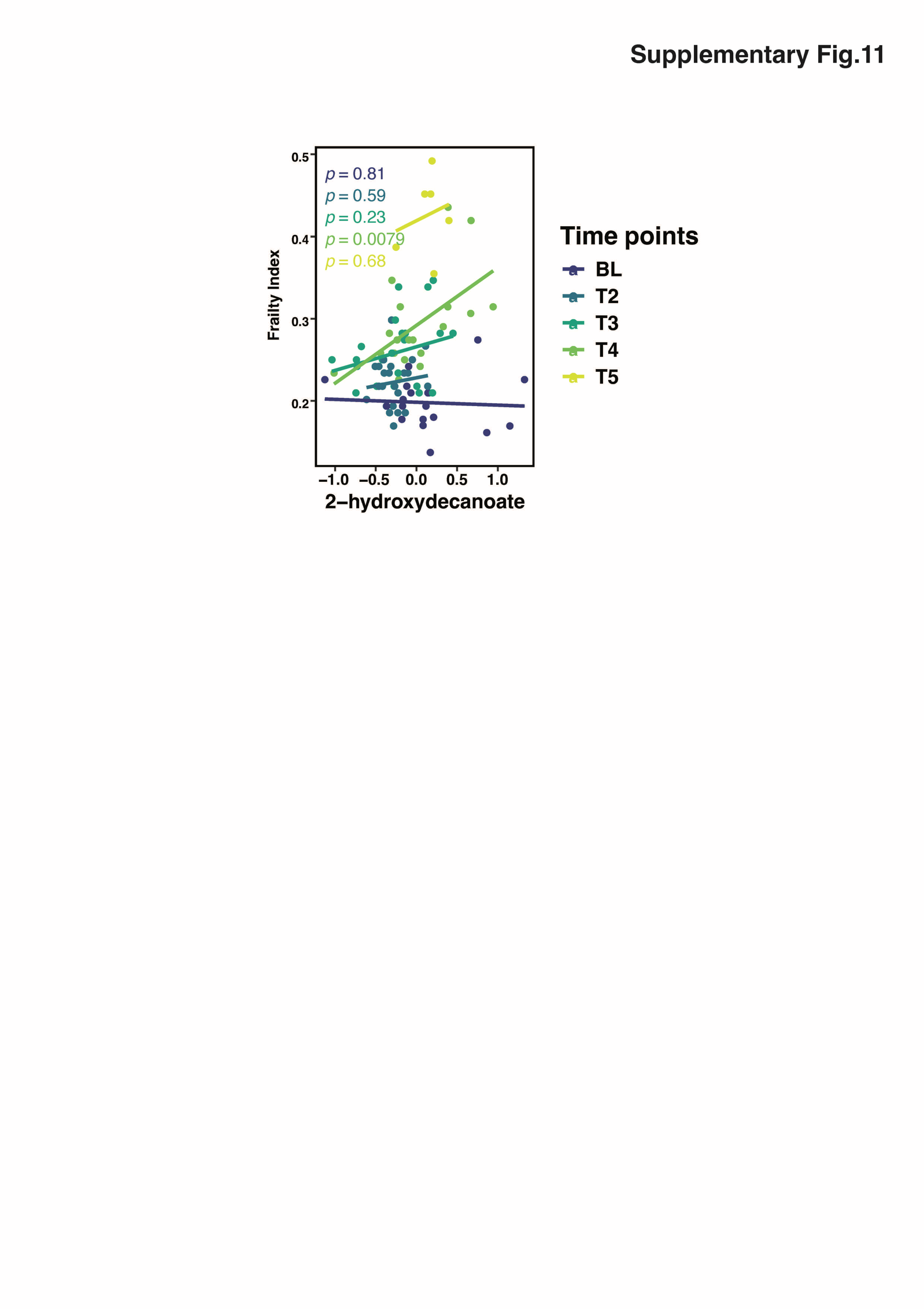

***Supplementary Fig.11. Association of 2-hydroxydecanoate with frailty index at five time points.***

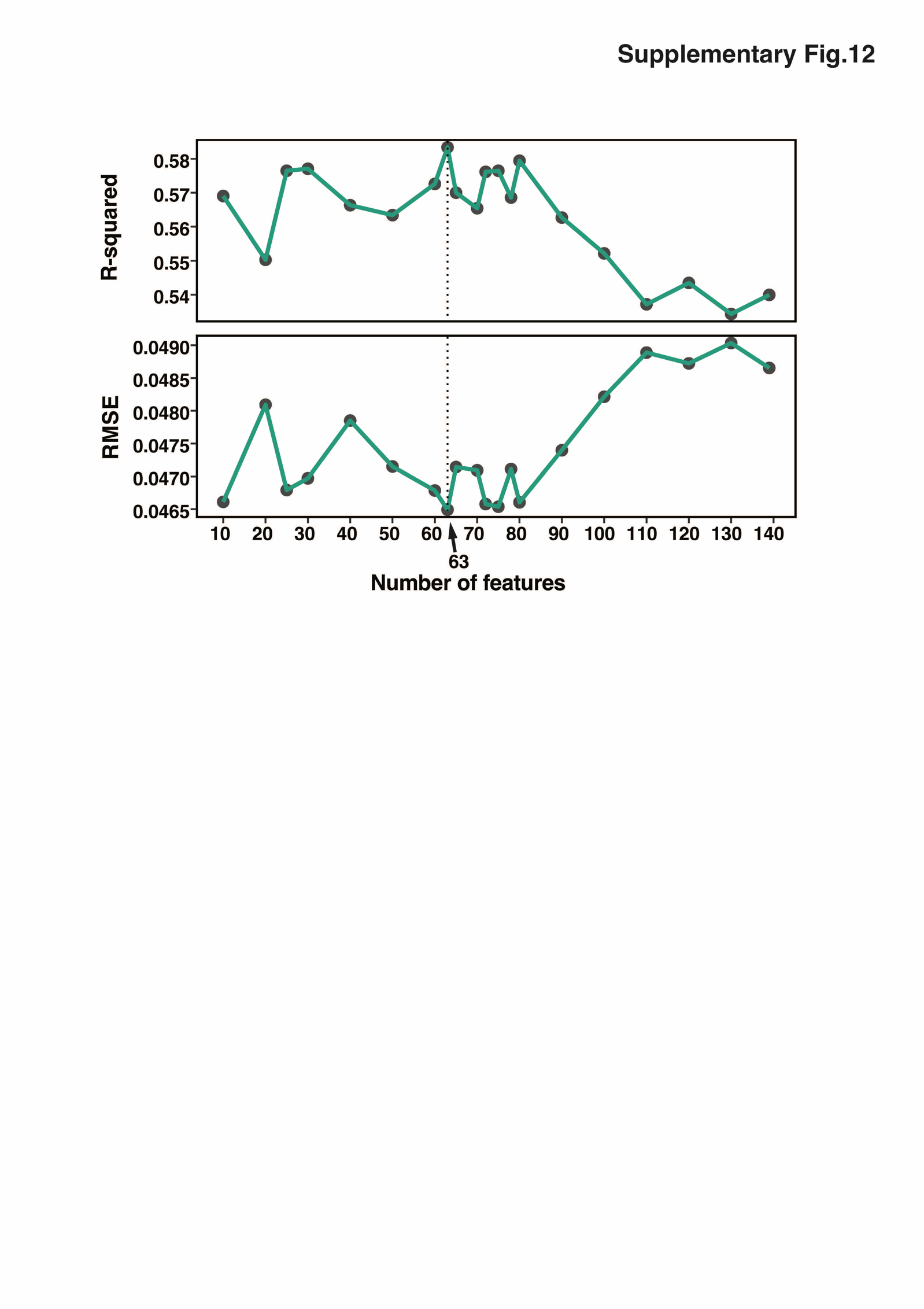

### Supplementary Fig.12. Model performance comparisons using different numbers of top informativity-based metabolite features.
